# Gut expressed vitellogenin is hijacked by a virus to facilitate its spread

**DOI:** 10.1101/2020.04.29.068825

**Authors:** Ya-Zhou He, Yu-Meng Wang, Tian-Yan Yin, Shu-Sheng Liu, Xiao-Wei Wang

## Abstract

Insect vitellogenin (Vg) has been considered to be synthesized in the female fat body, secreted into hemolymph, and taken up by developing oocytes to serve as nutrition for embryo development. However, the expression and function of Vg in other tissues are largely unknown. Here, we show that Vg is expressed in the whitefly midgut epithelial cells and secreted into gut lumen where it binds to the microvillar membrane. Furthermore, we find that the midgut expressed Vg is hijacked by *Tomato yellow leaf curl virus* to facilitate the movement of virus crossing the midgut wall. Silencing of Vg or immune-blocking the interaction between viral coat protein and midgut Vg inhibited virus movement across the midgut wall and decreased virus transmission. Our findings show possession of functional Vg in the midgut of an insect, and reveal a novel mechanism involved in the movement of virus crossing the midgut barrier of vector insect.

## Introduction

Vitellogenins (Vgs) are precursors of egg yolk that serves as the major source of nutrition for embryo development in almost all oviparous animals, including vertebrate and insect (1). Vgs are usually synthesized by extraovarian tissues, secreted into the circulatory system, and subsequently internalized by developing oocytes through receptor-mediated endocytosis (1, 2). Vertebrate Vgs are synthesized in the liver, and then secreted in the bloodstream (3). Insect Vgs are considered to be synthesized by the female fat body, an functional equivalent of vertebrate liver and adipose tissue, and then secreted into the hemolymph and taken up by developing oocytes (2). In recent years, accumulating data have shown that the female fat body is not the only the vitellogenic tissue of insect, as its synthesis, although in smaller quantities, also occurs in other female tissues as well as in males of some species (4). For example, Cyclorapha ovarian follicular cells produce Vgs (5). *Apis mellifera* Vg is synthesized in the ovaries, hypopharyngeal glands and the head fat bodies of functionally sterile honeybee workers (6). In *Bombus hypocrita*, the Vg gene is expressed at various levels in different castes, including the queen, workers and drones (7). Recently, Huo *et al*. report that Vg is sex-independently synthesized in the *Laodelphax striatellus* hemocytes (8). These results indicate that the function of insect Vg may extend beyond serving as nutrition for developing embryos.

Many viruses that cause diseases in humans, animals and plants are persistently transmitted by arthropod vectors, which notably encompass mosquitoes, ticks, whiteflies, leafhoppers, planthoppers and aphids (9, 10). After they are acquired from animal blood or plant sap by the arthropod, the virions must first overcome the gut infection and dissemination barriers to get into the hemolymph or other tissues, and finally into the salivary gland prior to being excreted and transmitted to new hosts (10). Previous studies showed that many persistent viruses can be transmitted at higher efficiency when they are experimentally delivered into the hemocoel of the vector than delivery via oral acquisition (11, 12), and that some viruses can even be transmitted by a non-vector when they are injected into the hemocoel of the insect (13, 14), illustrating the function of insect gut as a major barrier to persistent viruses transmission. Passage of viruses through the gut barrier requires specific interactions between virus and vector components (10, 15). Insect gut proteins that involved in virus movement across the gut wall are considered as important targets for virus transmission blockage (11, 16).

Begomoviruses (family *Geminiviridae*) are known as the largest genus of >400 species of plant viruses and are exclusively transmitted by whiteflies of the *Bemisia tabaci* cryptic species complex in a persistent manner (17, 18). In the past 30 years, two species of the *B. tabaci* complex, provisionally named as Middle East Asia Minor 1 (MEAM1, previously biotype B) and Mediterranean (MED, previously biotype Q), have invaded many regions of the world. Along with the invasion of these two species of whiteflies, begomoviruses have emerged worldwide as serious constraints to the cultivation of a variety of economically important crops (19, 20). Among these viruses, *Tomato yellow leaf curl virus* (TYLCV) is one of the most devastating viral agents and has caused extensive damage to many crops especially tomato (21, 22). TYLCV coat protein (CP) is considered the key viral component specifically interacting with whitefly proteins for virus transmission to occur (23, 24). Previous studies have provided evidence that TYLCV enters whitefly midgut epithelial cells through receptor-mediated, clathrin-dependent endocytosis, and that the early steps of endosome trafficking play an important role in the intracellular movement of the virus crossing the midgut wall (25, 26). However, whitefly proteins mediating these processes are not known.

In this study, using midgut-specific immunoprecipitation followed by high-throughput mass spectrometry proteomic analysis, we found that Vg was synthesized in the whitefly midgut and hijacked by TYLCV CP. Moreover, the interaction between viral CP and midgut Vg contributed to the movement of virus crossing the midgut wall for efficient transmission. Our results give valuable clues for the development of new strategies to block virus transmission, and provide novel insights into the function of insect Vg.

## Results

### Identification of whitefly proteins in the midgut that interact with TYLCV CP

To identify whitefly midgut proteins that interact with TYLCV CP, total proteins were isolated from 2000 midguts of viruliferous MEAM1 whiteflies and used for immunoprecipitation with a mouse anti-TYLCV CP monoclonal antibody or mouse pre-immune sera (control sera) (Fig. S1A). Western blotting assay showed that a single band at 30 kD (the predicted molecular weight of TYLCV CP) was detected in total proteins of viruliferous whiteflies, but not in that of non-viruliferous whiteflies (Fig. S1B), confirming the specificity of the anti-TYLCV CP antibody. Then, the immunoprecipitates were digested with trypsin and analyzed by shotgun ultra-performance liquid chromatography (UPLC)-tandem MS (MS/MS) to identify TYLCV CP binding partners in the midgut (Fig. S1A). Only proteins specifically co-immunoprecipitated by the anti-TYLCV CP antibody but not by the control sera and with at least two peptide-spectrum matches were considered credible candidates. Under these conditions, a total of 76 proteins were identified (Data S1). To our surprise, Vg protein was isolated from midgut proteins of viruliferous whiteflies through the above approach (Data S1).

### Vg is expressed in the midgut of whitefly

To validate the expression of Vg in the midgut, we first examined Vg transcript in the midgut and fat body of female MEAM1 whiteflies. Quantitative reverse transcription PCR (qRT-PCR) showed that both the midgut and fat body produced Vg mRNA, with higher levels in the fat body, and lower levels in the midgut. Moreover, whereas the mRNA level of Vg in the fat body dramatically increased with whitefly development, it was stable in the midgut of adult whiteflies at 1 day and 10 days after eclosion (DAE) (Fig. 1A). Immunoblotting analysis using an anti-Vg monoclonal antibody (Fig. S1C) (27) verified the existence of Vg protein in the midgut and fat body of whiteflies (Fig. 1 B). We further visualized Vg protein using immunofluorescence assays (IFA) with the anti-Vg monoclonal antibody. Specific signals were observed in the midgut and fat body of whiteflies at 1 and 10 DAE. (Fig. 1C). Consistent with the mRNA levels, the quantity of Vg protein was elevated in the fat body with whitefly development, but remained stable in the midgut (Fig. 1 B and C).

**Fig 1.**
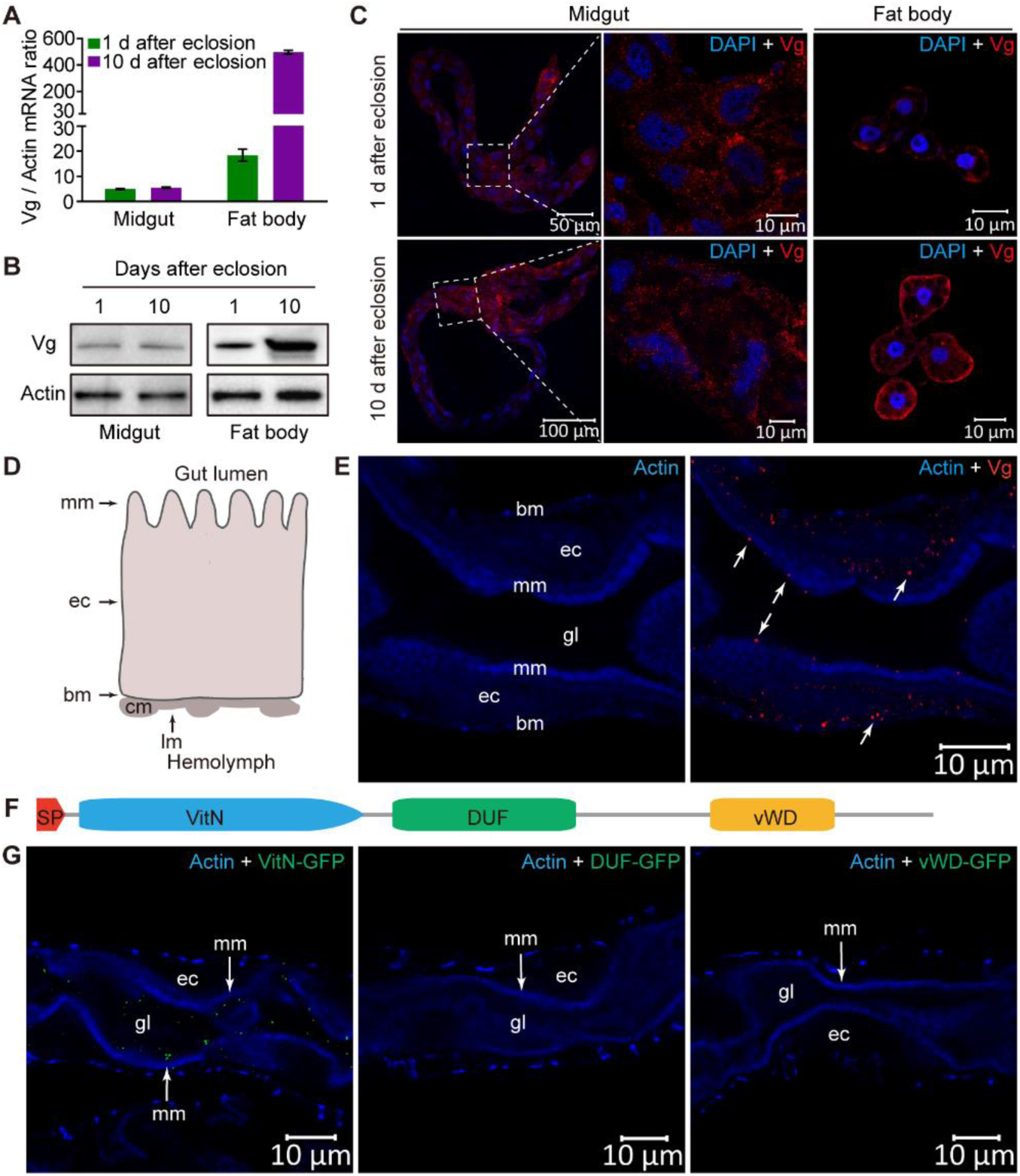
Vg is expressed in the midgut of whiteflies. (A and B) Vg mRNA (A) and protein (B) levels in the midgut and fat body of adult female whiteflies at two developmental stages. (A) Mean ± SEM from three independent experiments. (C) Immunofluorescence staining of Vg protein in midgut and fat body of adult female whiteflies at two developmental stages. Cell nucleus was stained with DAPI (blue). (D) Diagram of whitefly midgut epithelium. (E) Subcellular localization of Vg protein in midgut epithelial cells. The white arrow indicates the immune-reactive signal of Vg protein. (C and E) Vg was detected using a mouse anti-Vg monoclonal antibody and goat anti-mouse IgG labeled with Dylight 549 (red) secondary antibody. (F) Diagram illustrating the structural domains of whitefly full-length Vg protein. SP, signal peptide. VitN, vitellogenin N-terminal domain. DUF, domain of unknown function. vWD, von Willebrand factor type D domain. (G) The VitN domain of Vg functions in binding with the midgut microvillar membrane. VitN-GFP was detected using a rabbit anti-GFP monoclonal antibody and goat anti-rabbit IgG labeled with Dylight 488 (green) secondary antibody. (E and G) The actin-based microvillar membrane and muscle fibers were stained with Dylight 647 phalloidin (blue). *cm* circular muscle, *lm* longitudinal muscle, *gl* gut lumen, *mm* microvillar membrane, *ec* epithelial cell, *bm* basal membrane. Images are representative of three independent experiments with a total of 30 whiteflies analyzed for each treatment.

Insect Vgs are secreted proteins (4). To determine the distribution pattern of Vg in the midgut, we examined subcellular localization of Vg protein in midgut epithelial cells of female whiteflies. The alimentary canal of whitefly is composed of a single layer of epithelial cells, with microvillar membrane on the lumen side and basal membrane on the hemocoel side, covered with muscle fibers (Fig. 1D and Fig. S2A) (26). IFA showed the presence of Vg protein in the cytoplasm and at the lumen side of microvillar membrane (Fig. 1E), indicating that Vg is synthesized in the cytoplasm of midgut epithelial cells and secreted into gut lumen where it binds to the microvillar membrane. The whitefly Vg consists of three Vg-specific function domains: the vitellogenin N-terminal (VitN) domain, a middle-region domain of unknown function (DUF), and a von Willebrand factor type D (vWD) C-terminal domain (Fig. 1F) (27). To investigate whether Vg binds to the midgut microvillar membrane and which domains mediate the binding, the VitN-GFP, DUF-GFP and vWD-GFP fusions were expressed in *Drosophila Schneider* 2 (S2) cells (Fig. S3A) and used for *in vivo* midgut binding assays (28). Whiteflies were fed with recombinant proteins for 4 h followed by a 6 h feeding on a sucrose solution to remove unbound proteins. IFA showed that VitN-GFP could be readily detected at the lumen side of microvillar membrane. In contrast, neither DUF-GFP nor vWD-GFP was detected in whitefly midguts (Fig. 1G). Specificity of the binding was further confirmed by feeding whiteflies with similar amount of GFP and VitN-GFP (Fig. S3B), of which no binding was found for GFP alone (Fig. S3C). Overall, these results suggest that Vg is synthesized in the whitefly midgut epithelial cells and secreted into gut lumen where it binds to microvillar membrane through the VitN domain.

### TYLCV interacts with Vg in the whitefly midgut

To verify the interaction of TYLCV and Vg in the whitefly midgut, firstly, we examined the localization of TYLCV and Vg in the midgut of viruliferous whiteflies. The whitefly midgut consists of the gastric caecum, filter chamber, descending midgut and ascending midgut (Fig. S2B) (29). IFA showed that TYLCV co-localized with Vg in all these parts of the midgut (Fig. 2 A and B), confirming the association of TYLCV with Vg in the whitefly midgut. Next, we investigated whether Vg directly binds to TYLCV CP and which of the three Vg domains mediate the binding using an GST pull-down assay. The results showed that the VitN and vWD domains could directly bind to GST-fused TYLCV CP, but none bound to GST alone, and the VitN domain interacted more strongly with TYLCV CP than did vWD (Fig. 2A). These results, together with the previous immunoprecipitation followed by UPLC-MS/MS analyses (Data S1), demonstrate that TYLCV interacts directly with Vg in the whitefly midgut.

**Fig 2.**
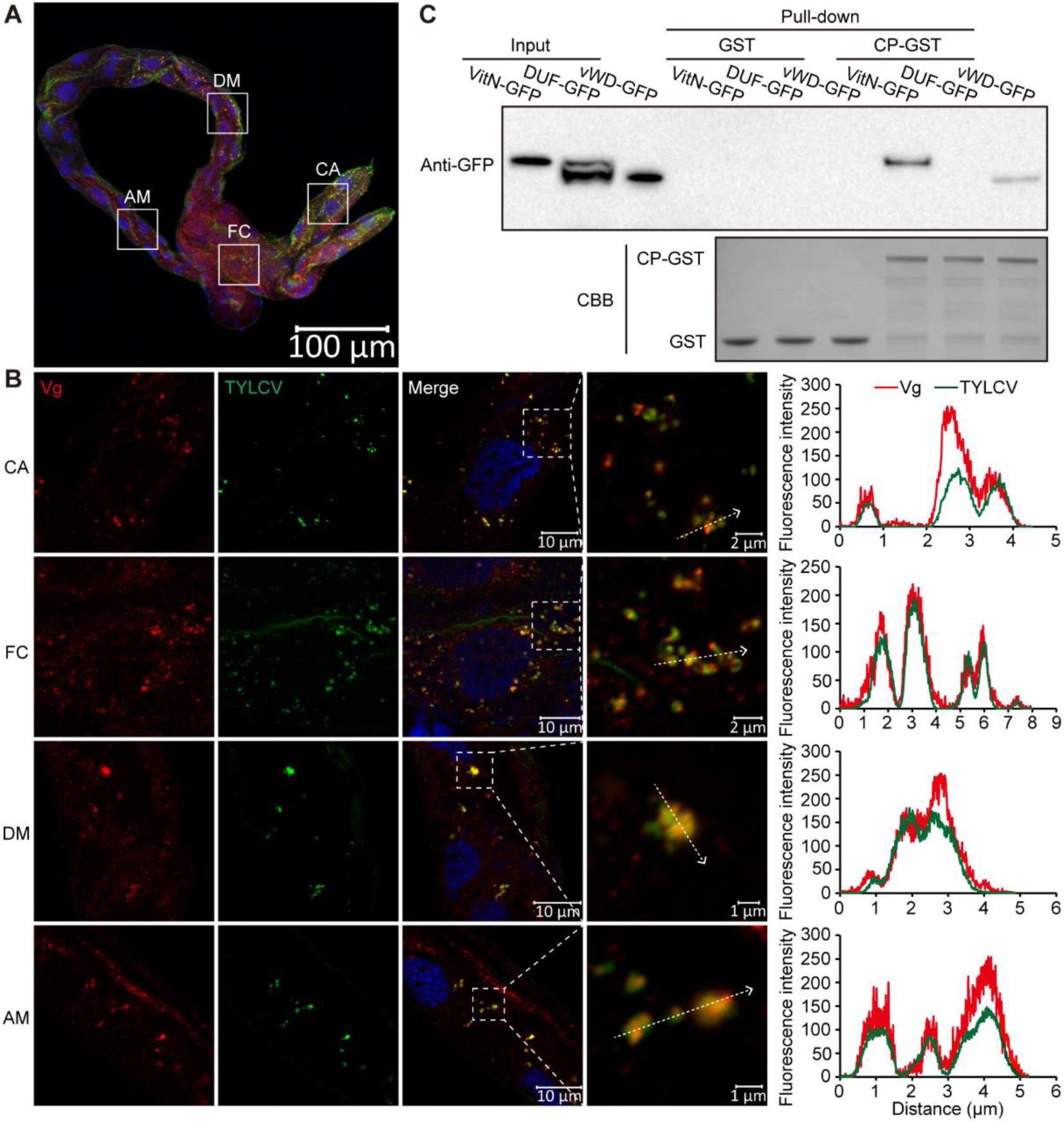
TYLCV interacts with Vg in the midgut. (A and B) Localization of Vg and TYLCV in different parts of the midgut from viruliferous female whitefly. Midguts of female whiteflies exposed to TYLCV-infected tomato plants for a 72 h acquisition access period (AAP) were dissected and used for immunofluorescence. Panels in (B) are the magnification of boxed area in gastric caecum (CA), filter chamber (FC), descending midgut (DM) and ascending midgut (AM) of the midgut in (A). Vg was detected using a mouse anti-Vg monoclonal antibody and goat anti-mouse IgG labeled with Dylight 549 (red) secondary antibody. TYLCV was detected using a rabbit anti-coat protein (CP) polyclonal antibody and goat anti-rabbit IgG labeled with Dylight 488 (green) secondary antibody. Cell nucleus was stained with DAPI (blue). Yellow color indicates the overlay of red and green. Analyses of overlapped fluorescence spectra from Vg and TYLCV in the boxed areas in (B) were shown on the right of the images. The white dashed arrows in (B) mark the line scans and the direction used to create the fluorescence intensity profiles. Images are representative of three independent experiments with a total of 30 whiteflies analyzed. (C) Mapping of Vg domains interact with TYLCV CP using GST pull-down assay. Coomassie Brilliant Blue (CBB) staining of purified GST and CP-GST serves as a loading control.

### TYLCV moves across the midgut epithelial cells as a complex with Vg

In order to understand the role of the interactions between TYLCV and Vg in virus movement across the midgut epithelial cells, firstly, we traced the infection process of TYLCV in the midgut by IFA. The infection process could be classified into five phases. In phase I, TYLCV virions were detected only in the filter chamber of 23% of the tested midguts of whiteflies after 1 h acquisition access period (AAP). In phase II, virions were detected in the gastric caecum and descending midgut of 40% of the tested midguts after 1 h AAP and 60% after 3 h AAP. In phase III, the virions were seen throughout the midgut in 44% of the tested midguts after 6 h AAP. In phase II and phase III, the virions were bound to the microvillar membrane and not seen in the cytoplasm. In phase IV, virions were seen in the cytoplasm of epithelial cells in 60% of the tested midguts after 12 h AAP. In phase V, virions were seen in the cytoplasm close to the basal membrane of epithelial cells in 50% of the tested midguts after 24 h AAP and 93% after 48 h AAP, respectively (Fig. S4 and Table S1). These observations showed that after entering insect midgut lumen with plant sap, TYLCV virions first bind to the microvillar membrane, then invade the cytoplasm and move to the basal membrane, where they can be released into the hemolymph for further spread.

Given that Vg protein is located at the lumen side of microvillar membrane and in the cytoplasm, we then examined where the TYLCV-Vg interaction occurred during virus movement across the midgut epithelial cells. IFA showed that TYLCV co-localized with Vg at the lumen side of microvillar membrane (Fig. 3A), in the microvillar membrane (Fig. 3B), in the middle of cytoplasm (Fig. 3C), and in place near to the basal membrane (Fig. 3D). suggesting that TYLCV binds to Vg at the lumen side of microvillar membrane and then moves first to the cytoplasm and then to the basal membrane as a complex with Vg.

**Fig 3.**
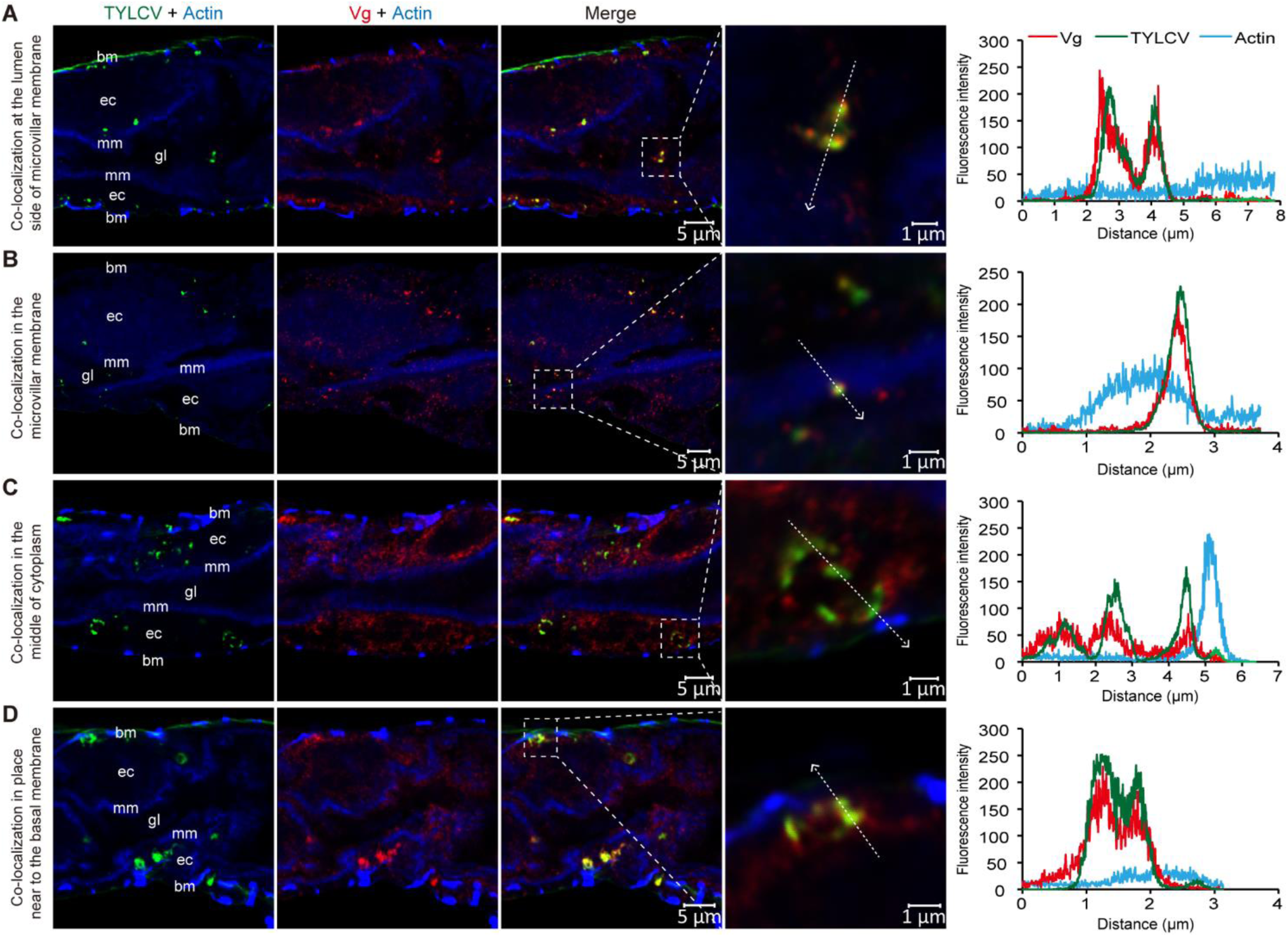
Subcellular localization of TYLCV and Vg in midgut epithelial cells. TYLCV co-localized with Vg at the lumen side of microvillar membrane (A), in the microvillar membrane (B), in the middle of cytoplasm (C) and in place near to the basal membrane (D). Midguts of female whiteflies exposed to TYLCV-infected plants for a 72 h AAP were dissected and used for immunofluorescence. TYLCV was detected using a rabbit anti-CP polyclonal antibody and goat anti-rabbit IgG labeled with Dylight 488 (green) secondary antibody. Vg was detected using a mouse anti-Vg monoclonal antibody and goat anti-mouse IgG labeled with Dylight 549 (red) secondary antibody. Yellow color indicates co-localization of red and green. Actin-based microvillar membrane and visceral muscles were stained with Dylight 647 phalloidin (blue). Analyses of fluorescence spectra from Vg, TYLCV and Actin in the boxed area were shown on the right of the images. The white dashed arrows mark the line scans and the direction used to create the fluorescence intensity profiles. Images are representatives of multiple experiments with multiple preparations. *gl* gut lumen, *mm* microvillar membrane, *ec* epithelial cell, *bm* basal membrane.

Previous studies have shown that vesicle trafficking is critical for the movement of TYLCV across the midgut wall in whiteflies. Especially, almost all TYLCV co-localized with *Helix pomatia* agglutinin (HPA), a *N*-acetylgalactosamine binding lectin (30), labelled vesicles within midgut epithelial cells (26). The above results indicate that TYLCV binds to Vg and moves across the midgut wall as a complex with Vg. If this hypothesis were correct, then Vg protein would also co-localized with HPA-labelled vesicles within midgut epithelial cells. Consequently, we examined the localization of TYLCV, Vg and HPA-labelled vesicles in the midgut of viruliferous whiteflies, and found that both TYLCV and Vg co-localized with HPA-labelled vesicles in the midgut (Fig. 4A and Fig. S5). It has been found later that the early endosomes are essential for the movement of TYLCV across the whitefly midgut wall. TYLCV was found to be co-localized with early endosomes in the midgut epithelial cells (26). To further confirm the association of TYLCV and Vg during the intracellular movement of TYLCV, we examined whether TYLCV and Vg co-localize with early endosomes using a specific antibody against Rab5 (26), an early endosome marker protein. The results showed that both TYLCV and Vg co-localized with Rab5-labelled early endosomes within midgut epithelial cells (Fig. 4B). Taken together, these results clearly show that TYLCV moves across the midgut epithelial cells as a complex with Vg, suggesting an important role of the midgut expressed Vg in virus movement across the midgut wall.

**Fig 4.**
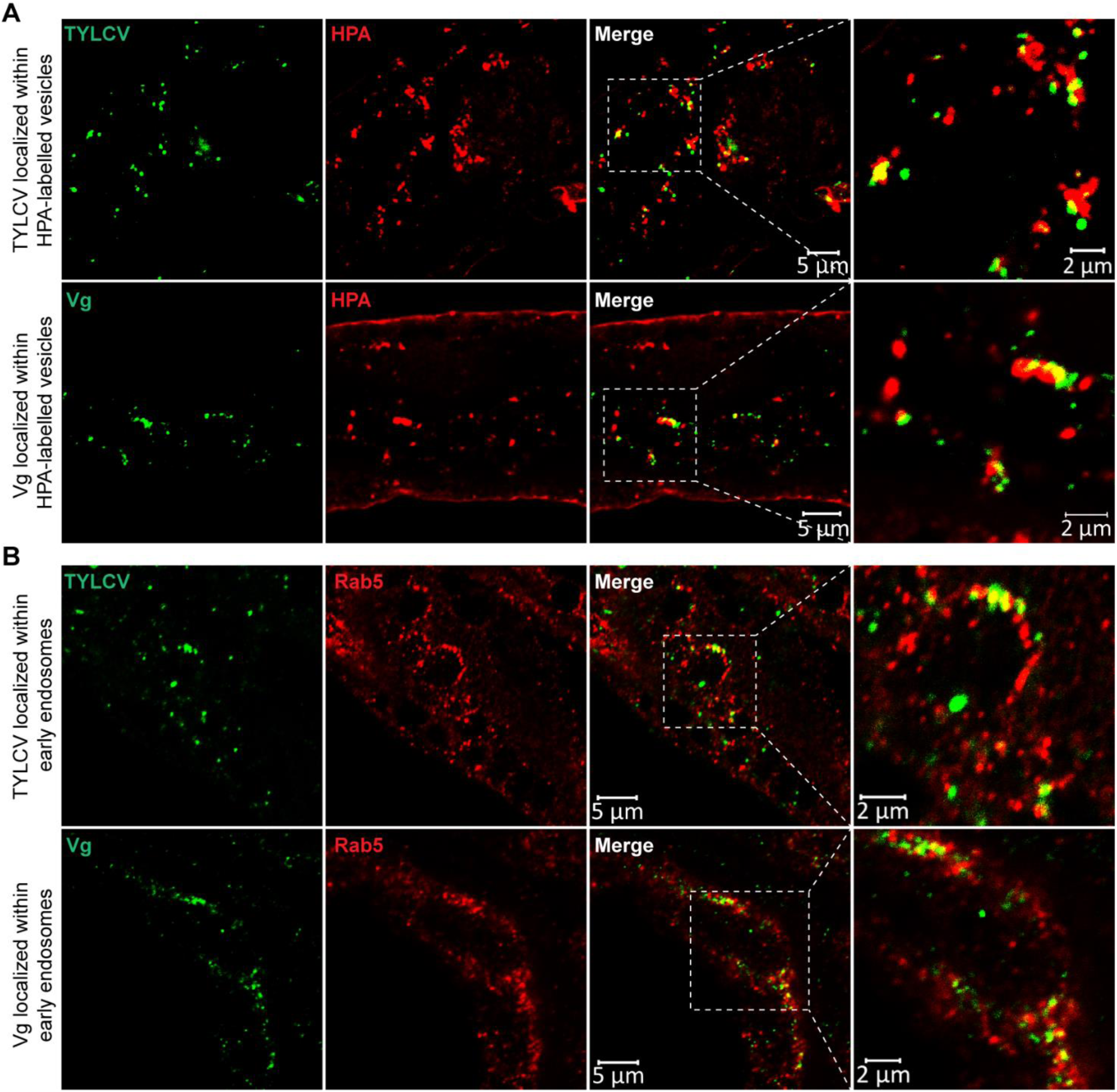
Co-localization of TYLCV and Vg with intracellular vesicles and early endosomes. Both TYLCV and Vg co-localized with lectin HPA-labelled vesicles (A) and Rab5-labelled early endosomes (B). Midguts of female whiteflies exposed to TYLCV-infected plants for a 72 h AAP were dissected and used for immunofluorescence. TYLCV was detected using a rabbit anti-CP polyclonal antibody and goat anti-rabbit IgG labelled with Dylight 488 (green) secondary antibody. Vg was detected using a mouse anti-Vg monoclonal antibody and goat anti-mouse IgG labelled with Dylight 488 (green) secondary antibody. Intracellular vesicles were labelled using Alex 647 lectin HPA (red). Early endosomes were detected using a rabbit anti-Rab5 polyclonal antibody and goat anti-rabbit IgG labelled with Dylight 549 (red) secondary antibody. Yellow color indicates co- localization of red and green. Images are representative of multiple experiments with multiple preparations.

### Silencing Vg inhibits TYLCV movement across the midgut wall

To examine the role of Vg in the movement of TYLCV across the midgut wall, we knocked down the expression of Vg using RNA interference (RNAi). Compared with whiteflies fed with dsGFP (control), the Vg mRNA level in whitefly whole body decreased by 55% in the group fed with dsVg and Vg protein level also decreased (Fig. 5A). The Vg expression level in the midgut of dsVg-treated whiteflies also exhibited an obvious reduction as shown by IFA analysis (Fig. S6). In order to compare the virus acquisition efficiency, we put two groups of whiteflies on two opposite leaflets of a TYLCV-infected tomato plant (Fig. S7A). Quantitative PCR (qPCR) showed that, without treatment, quantities of virus acquired by whiteflies after a 24 h AAP on two opposite leaflets were similar (Fig. S7B). In dsVg-treated whitefly whole bodies, the abundance of TYLCV DNA decreased by 58% after 24 h AAP on TYLCV-infected plants compared with that of the dsGFP-treated control group (Fig. 5B). IFA further showed that TYLCV had invaded the cytoplasm of the majority of midgut epithelial cells of dsGFP-treated whiteflies (Fig. 5C), and the proportion of midguts in phase V in the dsVg treatment (26%) was apparently lower than the dsGFP control (51%) (Table S2), indicating that dsVg treatment decelerated TYLCV infection of the midgut.

**Fig 5.**
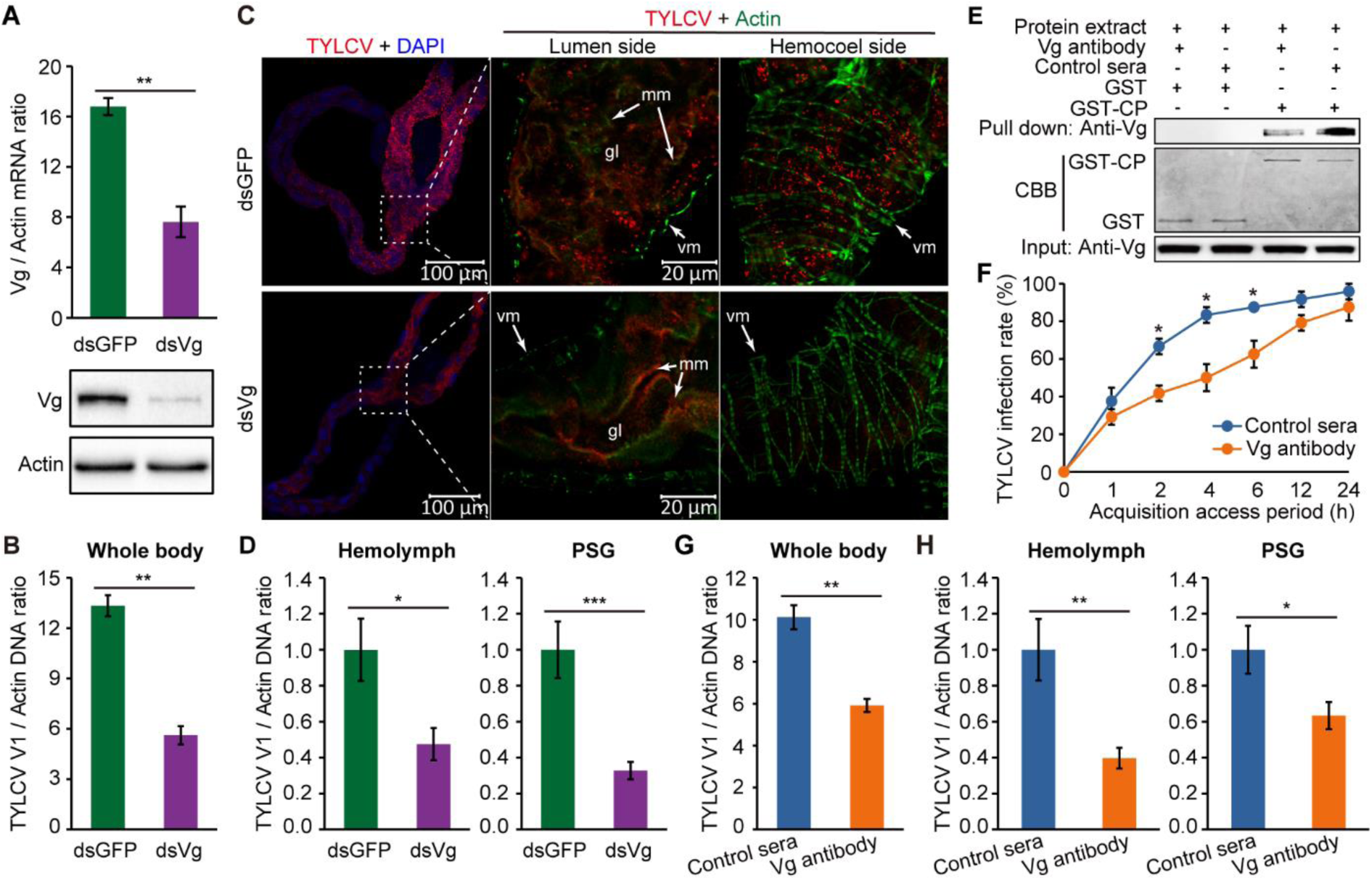
The role of Vg in TYLCV movement across the midgut wall. (A) Vg mRNA and protein levels in whitefly whole body after feeding with dsRNAs. (B) TYLCV DNA levels in whitefly whole body after a 24 h AAP on TYLCV-infected tomato plants following dsRNAs treatment. (C) Localization of TYLCV in midguts of dsRNAs-treated whiteflies after a 24 h AAP on TYLCV-infected plants. TYLCV was detected using a mouse anti-CP monoclonal antibody and goat anti-mouse IgG labelled with Dylight 549 (red) secondary antibody. Cell nucleus was stained with DAPI (blue). Actin-based microvillar membrane and visceral muscles were stained with Dylight 647 phalloidin (green). *gl* gut lumen, *mm* microvillar membrane, *vm* visceral muscles. Images are representative of three independent experiments with a total of 60 whiteflies analyzed for each treatment. (D) TYLCV DNA levels in the hemolymph and primary salivary gland (PSG) of dsRNAs- treated whiteflies after a 24 h AAP on TYLCV-infected plants. (E) Vg antibody interfered with the interaction between Vg and TYLCV CP. CBB staining of purified GST and CP-GST serves as a loading control. (F) TYLCV infection rate of Vg antibody or mouse pre-immune (control sera)- treated whiteflies after various periods of time from 0 to 24 h feeding on TYLCV-infected plants. Each time point (n=24). (G) TYLCV DNA levels in Vg antibody or control sera-treated whitefly whole bodies after a 24 h AAP on TYLCV-infected plants. (H) TYLCV DNA levels in the hemolymph and PSG of Vg antibody or control sera-treated whiteflies after a 24 h AAP on TYLCV-infected plants. (A-B and F-G) Mean ± SEM from three independent experiments. ^*^, *p* < 0.05, ^**^, *p* < 0.01 (independent-samples t-test). (D and H) Mean ± SEM from 29-30 independent samples. One PSG or the hemolymph of one female whitefly was used as one sample for analyzing virus quantity. ^*^, *p* < 0.05, ^**^, *p* < 0.01, ^***^, *p* < 0.001 (non-parametric Mann-Whitney U test).

Next, the quantity of TYLCV that had moved across the midgut epithelial cells was examined by quantifying virus abundance. Compared with the control, the abundance of TYLCV DNA in the hemolymph and primary salivary glands (PSGs) of dsVg-treated whiteflies decreased by 52% and 75% respectively (Fig. 5D). In a virus infection assay, 17% and 30% of the plants in the dsVg treatment were symptomatic at 15 d and 30 d post-transmission, while in the control 37% and 43% of the plants were symptomatic at 15 d and 30 d post-transmission, indicating that interference with Vg reduced virus transmission and infection (Table S2). Taken together, these results indicate that the reduction in Vg expression by RNAi significantly inhibited the movement of TYLCV across the midgut wall, leading to reduction of transmission efficiency of the whitefly vector.

### Disrupting the interaction between viral CP and midgut Vg inhibits TYLCV movement across the midgut wall

We further examined the function of the interaction between TYLCV CP and midgut Vg in the movement of TYLCV across the midgut wall by immune-blocking experiments using the mouse anti-Vg antibody. *In vitro* GST-pull down assay showed that the Vg antibody interfered with Vg binding to GST-fused TYLCV CP (Fig. 5E). After oral ingestion of Vg antibody, the midgut, fat body and ovary of female whiteflies were dissected for Vg antibody detection. IFA showed abundant Vg antibodies in the midguts but no Vg antibodies in the fat body or ovary. Meanwhile, no specific signal was detected in the midguts of mouse pre-immune serum (control sera)-fed whiteflies (Fig. S8A). These results indicate that the ingested Vg antibody mainly located in the midgut of whitefly through binding with midgut Vg. Therefore, oral ingestion of Vg antibody inhibits TYLCV CP-Vg interaction in the midgut and thus could be used to learn specifically the role of midgut Vg in the movement of TYLCV across the midgut wall.

We first examined the virus acquisition efficiency of Vg antibody-treated female whiteflies during a 24 h feeding on TYLCV-infected plants. Compared with whiteflies fed with control sera, the Vg antibody-fed whiteflies had significantly lower proportion of viral infection from 1-6 h (Fig. 5F). Although nearly all of the insects in each of the two treatments were infected with TYLCV after 24 h, the abundance of viral DNA in the whole body of Vg antibody-fed whiteflies were significantly lower than the control (Fig. 5 F and G). We next investigated the impact of Vg antibody treatment on the movement of TYLCV across the midgut wall. After a 24 h AAP, the proportion of midguts in phase V in the Vg antibody treatment (30%) was apparently lower than the control (55%) (Fig. S8B and Table S2), and the abundance of TYLCV DNA in the hemolymph and PSGs of Vg antibody-treated whiteflies was significantly lower than the control (Fig. 5H). In a virus infection assay, the proportions of symptomatic plants at 15 d and 30 d post-transmission in the Vg antibody treatment were reduced by 27% and 35% compared with the control (Table S2). Overall, the three sets of data indicate, similar to the effect of dsVg treatment, the immune-blocking of midgut Vg likewise reduced virus acquisition and transmission efficiency of the whitefly vector.

### Vg is expressed in the midgut of male whiteflies and involved in TYLCV transmission

Several studies have reported that insect Vg is also synthesized in males of some species, including the *Leucophaea maderae, Bombus hypocrite, Nicrophorus vespilloides, Camponotus festinatus* and *Laodelphax striatellus* (7, 8, 31-33). We thus investigated whether Vg is also expressed in the midgut of male MEAM1 whiteflies. qRT-PCR showed that Vg mRNA was produced in male midguts, with dramatically lower level than that of the females (Fig. 6A). IFA further confirmed the existence of Vg protein in midgut epithelial cells of males (Fig. 6B). To examine whether the Vg protein in male midgut was also involved in TYLCV transmission, we localized TYLCV and Vg in midgut epithelial cells of viruliferous males and found that TYLCV also co-localized with Vg (Fig. 6C). Next, we knocked down the expression of Vg in males using RNAi, and the Vg expression level in whole body was reduced by 72% in dsVg-treated males compared with the control (Fig. 6D). The virus quantity in dsVg-treated male whole bodies was significantly lower than that of the control after a 24 h AAP (Fig. 6E), suggesting that Vg also facilitates the movement of TYLCV across midgut wall in males.

**Fig 6.**
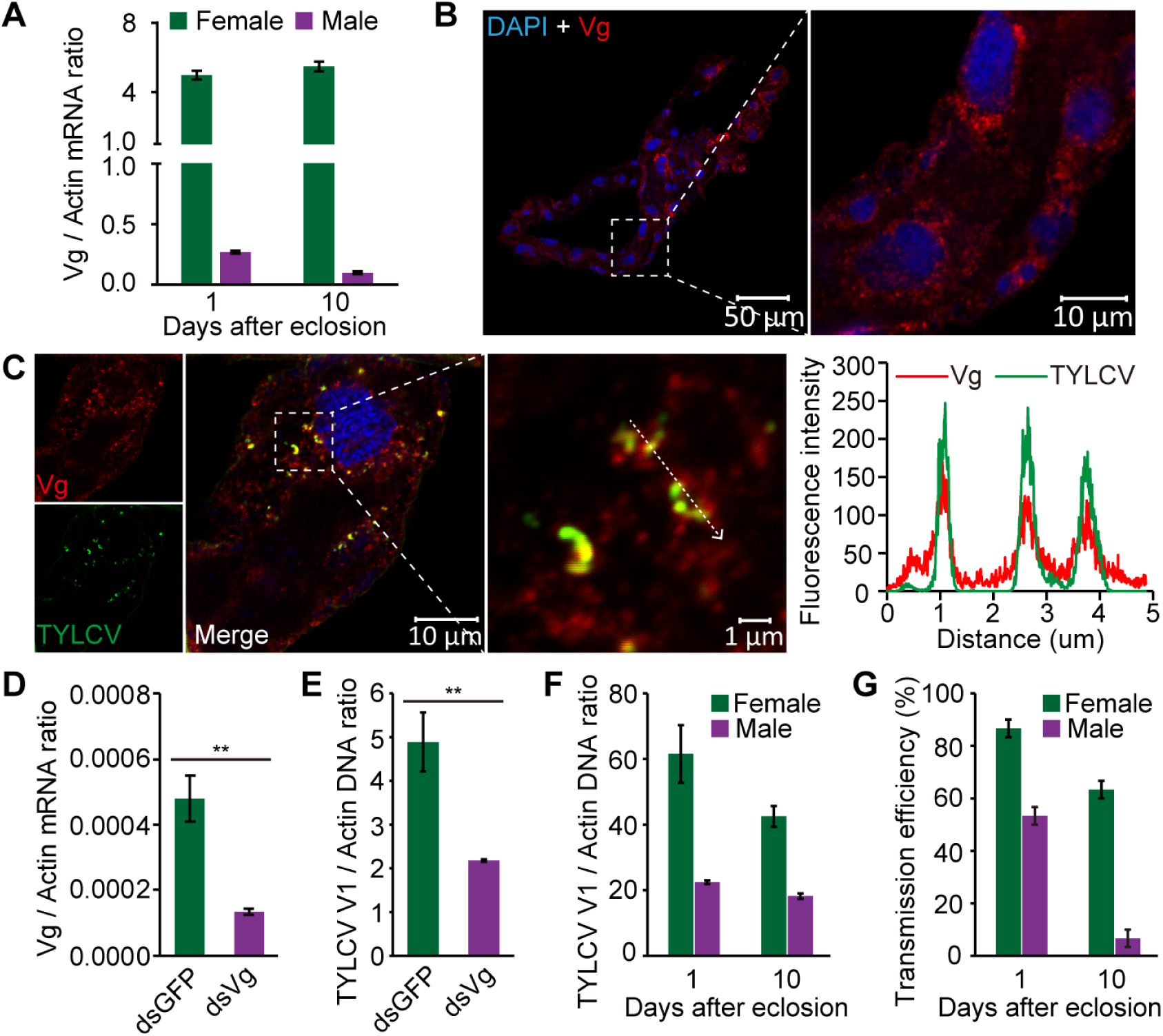
The role of Vg in TYLCV movement in male whiteflies. (A) Vg mRNA levels in the midgut of adult female and male MEAM1 whiteflies at two developmental stages. (B) Immunostaining of Vg protein in the midgut of male whiteflies at 1 day after eclosion. (C) Localization of Vg and TYLCV in midgut epithelial cells of viruliferous male whiteflies. Vg was detected using a mouse anti-Vg monoclonal antibody and goat anti-mouse IgG labelled with Dylight 549 (red) secondary antibody. TYLCV was detected using a rabbit anti-CP polyclonal antibody and goat anti-rabbit IgG labelled with Dylight 488 (green) secondary antibody. Cell nucleus was stained with DAPI (blue). Yellow color indicates co-localization of red and green. Analysis of overlapped fluorescence spectra from Vg and TYLCV in the boxed area was shown on the right of the image. The white dashed arrow marks the line scans and the direction used to create the fluorescence intensity profiles. Images are representative of multiple experiments with multiple preparations. (D) Vg mRNA levels in whole body of male whiteflies after feeding with dsRNAs. (E) TYLCV DNA levels in whole body of dsRNAs-treated males after a 24 h AAP on TYLCV-infected plants. (F) TYLCV DNA levels in whole body of female and male whiteflies at two developmental stages after a 48 h AAP on TYLCV-infected plants. (G) The inoculation capability of female and male adults at two developmental stages after a 48 h AAP on TYLCV-infected plants. For each combination, ten plants per replicate and three replicates were conducted to examine the transmission efficiency. (A and D-G) Mean ± SEM of three independent experiments. (D and E) ^**^, *p* < 0.01 (independent-samples t-test).

Given the different expression levels of Vg in the midgut of female and male whiteflies (Fig. 6A), we further compared the viral acquisition and transmission efficiencies between the two sexes. The results showed that the viral acquisition and transmission efficiency was positively correlated with Vg expression levels in the midgut of female and male whiteflies (Fig. 6 F and G). Therefore, the different expression levels of Vg in the midgut of female and male whiteflies may explain in part the differential acquisition and transmission efficiency of TYLCV by the two sexes, although there are also other reasons as reported by previous studies (34).

### The role of TYLCV-Vg interaction is conserved among different species of the *B. tabaci* complex

Previous studies have showed that TYLCV is also efficiently acquired by the MED and Asia II 1 cryptic species of the *B. tabaci* complex (35), we thus investigated whether the TYLCV-Vg interaction also exists in the midgut of these two whitefly species. qRT-PCR showed that Vg was expressed in the midgut of MED and Asia II 1 whiteflies (Fig. 7A). Similar to MEAM1 Vg, Vg proteins of both MED and Asia II 1 specifically interacted with GST-fused TYLCV CP but not with GST alone (Fig. 7B), suggesting a conserved interaction between TYLCV CP and *B. tabaci* Vg. IFA further showed the interaction of TYLCV and Vg in the midgut epithelial cells of these two whitefly species (Fig. 7 C and D). We then knocked down the expression of Vg in these two species using RNAi. Compared with the control group, the Vg expression levels in whitefly whole body decreased by 48% and 54% in dsVg-treated MED and Asia II 1 whiteflies, respectively (Fig. 7E). After a 24 h AAP on TYLCV-infected plants, the TYLCV abundance was reduced in both the dsVg-treated MED and Asia II 1 whiteflies compared with the control (Fig. 7F). Moreover, disrupting the interaction between viral CP and midgut Vg using the anti-Vg antibody also significantly decreased the TYLCV quantity in whiteflies after a 24 h AAP (Fig. 7G). Together, these data demonstrate that the role of TYLCV-Vg interaction in facilitating the movement of TYLCV across the midgut wall is conserved among different species of the *B. tabaci* complex.

**Fig 7.**
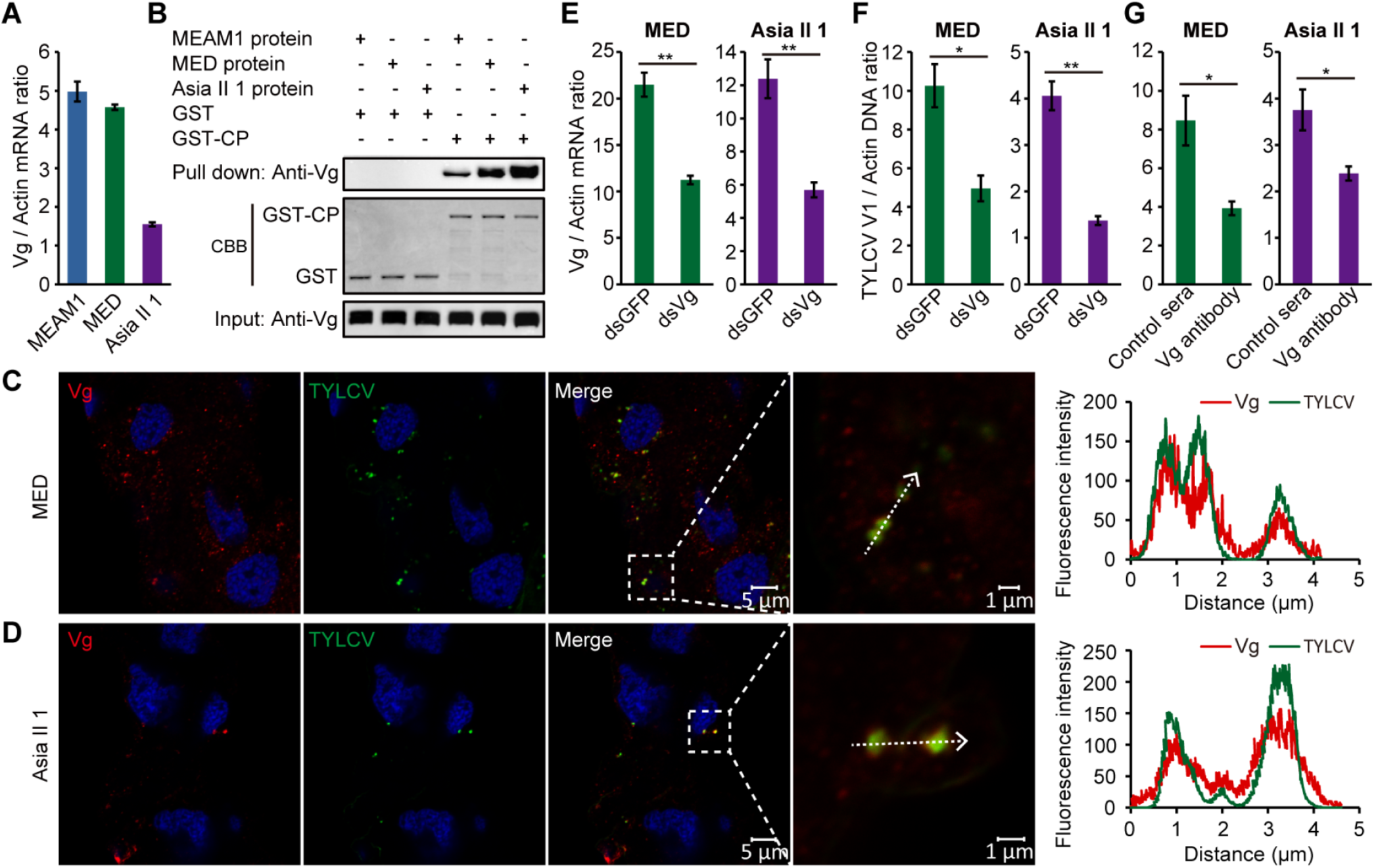
TYLCV-Vg interaction is conserved in different whitefly species. (A) Vg mRNA levels in midguts of different whitefly species. Total RNA was extracted from midguts of female MEAM1, MED or Asia II 1 whiteflies 1-2 days post-emergence, respectively, and assessed using qRT-PCR. Mean ± SEM of three independent experiments. (B) Vgs from three whitely species were all co- eluted with GST-fused CP but not with GST alone. (C and D) Localization of Vg and TYLCV in midgut epithelial cells of viruliferous whiteflies of MED (C) and Asia II 1 (D). Vg was detected using a mouse anti-Vg monoclonal antibody and goat anti-mouse IgG labeled with Dylight 549 (red) secondary antibody. TYLCV was detected using a rabbit anti-CP polyclonal antibody and goat anti- rabbit IgG labeled with Dylight 488 (green) secondary antibody. Cell nucleus was stained with DAPI (blue). Yellow color indicates the overlay of red and green. Analyses of overlapped fluorescence spectra from Vg and TYLCV in the boxed area were shown on the right of the images. The white dashed arrows mark the line scans and the direction used to create the fluorescence intensity profiles. Images are representative of three independent experiments with a total of 30 whiteflies analyzed for each treatment. (E) Vg mRNA levels in whitefly whole body after feeding with dsRNAs. (F) TYLCV DNA levels in whitefly whole body after a 24 h AAP on TYLCV-infected tomato plants following dsRNAs treatment. (G) TYLCV DNA levels in whitefly whole body after a 24 h AAP on TYLCV- infected tomato plants following anti-Vg antibody or control sera treatment. (E-G) Mean ± SEM of three independent experiments. ^*^, *p* < 0.05, ^**^, *p* < 0.01 (independent-samples t-test).

## Discussion

Insect Vgs are considered to be synthesized mainly by the female fat body (4). Nevertheless, it is clear now that Vg synthesis also occurs in other female tissues, such as ovaries, hypopharyngeal glands and hemocytes, as well as in males of some species of insects (5-8, 31-33). In the present study, the expression of Vg in the whitefly midgut was confirmed in several ways: (*i*) Vg transcript was detected in the midgut by qRT-PCR, and Vg protein was verified by western blotting and IFA using an anti-Vg monoclonal antibody (Fig. 1A-C); (*ii*) whereas the expression of Vg in the fat body increased with whitefly development, it remained stable in the midgut, eliminating possible contamination from fat body during dissection; (*iii*) IFA showed the presence of Vg protein in the cytoplasm and at the lumen side of microvillar membrane, indicating that Vg is synthesized in the cytoplasm of midgut epithelial cells and secreted into gut lumen where it binds to the microvillar membrane. The microvillar membrane binding ability of Vg was confirmed by *in vivo* midgut binding assays (Fig. 1D-G); and (*iv*) Vg is also expressed in the midgut of male whiteflies (Fig. 6). Thus, our findings expand Vg synthesis to a new important tissue-the insect midgut, a tissue absorbs the nutrients necessary for insect survival (36). Given the vital role of Vg in the development of all oviparous species, whether Vg is also expressed in the gut of other oviparous animals warrants further investigations.

Insect Vgs are secreted proteins. After synthesis in the fat body, Vgs are secreted into the hemolymph and then absorbed into growing oocytes via receptor-mediated endocytosis (4). Patterns of subcellular distribution of Vg protein in the midgut epithelial cells suggest that after synthesis in the cytoplasm Vg is secreted into the gut lumen, where it binds to the microvillar membrane (Fig. 8). Our *in vivo* midgut binding assays using three recombinant Vg domains showed that the N-terminal VitN domain of Vg bound to midgut microvillar membrane (Fig. 1D-G). Previous studies have shown that the N-terminal region of Vg is required for interacting with Vg receptor in the tilapia *Oreochromis aureus* and freshwater prawn *Macrobrachium rosenbergii* (37, 38). Therefore, the N- terminal region of Vg is likely responsible for receptor recognition in both vertebrate and invertebrate, including insects. It has been shown that in addition to supplying developing embryos with amino acids, Vgs are extensively modified, carrying covalently linked carbohydrates, phosphates and sulfates, and noncovalently bound lipids, vitamins, hormones and metals, thus facilitating the movement of these nutrients from hemolymph to ovaries (39). We thus speculate that the midgut expressed Vg may function in mediating the absorption of nutrients from the gut lumen into epithelial cells and finally into the hemolymph.

**Fig 8.**
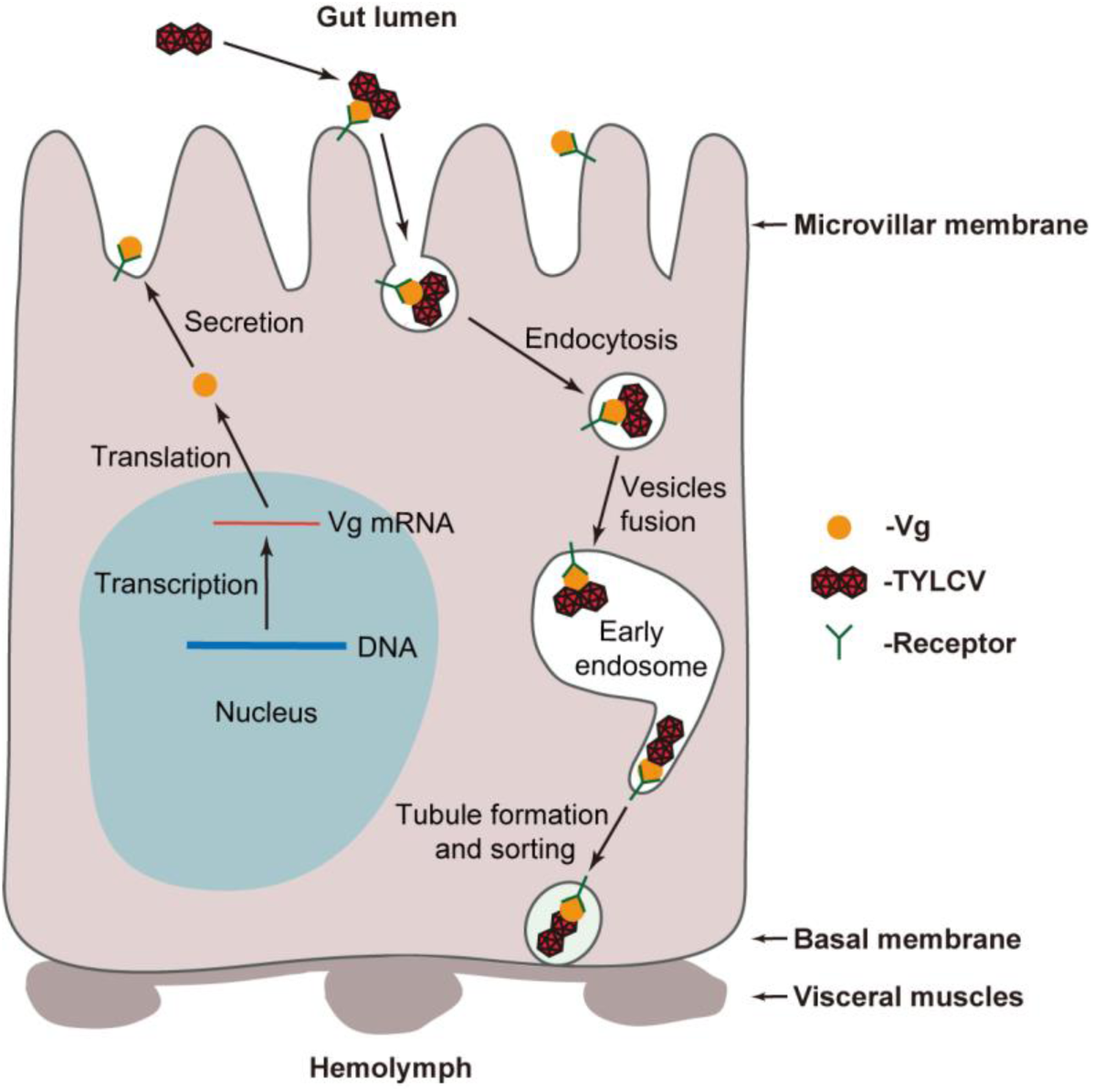
Model depicting Vg synthesis and hijacking of Vg by TYLCV in the midgut epithelial cell. Vg is synthesized in the cytoplasm of midgut epithelial cell and secreted into gut lumen where it binds to the microvillar membrane. After entering the midgut lumen with plant sap, TYLCV binds to Vg at the microvillar membrane and then entry into midgut epithelial cell through receptor-mediated, clathrin-dependent endocytosis. The virus-containing vesicle then deliver the virus-Vg complex to early endosome for further intracellular movement.

An essential step in the life cycle of many viruses is transmission to a new host by arthropod vectors, and one critical step in the transmission of persistently transmitted viruses is overcoming the midgut barrier to enter into vectors (9, 40). The insect midgut microvillar membrane acts as a barrier for viral entry, and the basal membrane as a barrier for viral escape (41, 42). Most viruses infect vector midgut epithelial cells via specific interaction between viral structural proteins and vector cell surface receptor complexes (16, 41, 43). Our results showed that the midgut expressed Vg was hijacked by a virus when moving across the midgut epithelial cells, starting at the lumen side of microvillar membrane, passing through the cytoplasm, and ending in places close to the basal membrane (Fig. 2, Fig. 3 and Fig. 4), suggesting that the midgut Vg is involved in the cell entry and intracellular movement of the virus in vector midgut. When knocking down the expression of Vg using RNAi or disrupting the interaction between viral CP and Vg in the midgut using anti-Vg antibody, the movement of virus across the midgut wall was inhibited and the virus transmission efficiency decreased (Fig. 5 and Table S2). Therefore, the midgut expressed Vg acts as an insect midgut receptor of virus. These findings not only improve understanding on virus-vector interactions but also provide clues for developing novel technology for blocking virus transmission.

The uptake of Vg by membrane-bound receptors (VgRs) occurs via receptor-mediated endocytosis (44). After binding, Vg/VgR complexes concentrate in clathrin-coated pits that invaginate and pinch off to form intracellular coated vesicles. These vesicles then carry the Vg to an endosome, which directs subsequent intracellular movement of Vg (1, 2). Not coincidentally, previous studies have shown that TYLCV enters whitefly midgut epithelial cells through receptor-mediated, clathrin-dependent endocytosis and that the early steps of endosome trafficking facilitate intracellular movement of TYLCV in whitefly midgut (25, 26). Our results show that both TYLCV and Vg co- localized with lectin HPA-labelled vesicles or Rab5-labelled early endosomes in the midgut epithelial cells (Fig. 4 and Fig. S5). Therefore, after entering the midgut lumen with plant sap, TYLCV may bind to Vg at the microvillar membrane and then entry into midgut epithelial cells through receptor-mediated, clathrin-dependent endocytosis. The virus-containing vesicles then deliver the virus-Vg complexes to early endosomes for further intracellular movement (Fig. 8).

Consistent with the evolutionarily conserved biochemical properties of insect Vgs (4), the TYLCV CP-Vg interaction was observed in several species of the *B. tabaci* complex. Moreover, this interaction is required for the movement of TYLCV across the midgut wall of all three whitefly species examined in this study, exhibiting a conserved mechanism used by TYLCV to overcome the midgut barrier of different species of insect vectors (Fig. 7). As the >400 species of begomoviruses are believed to be exclusively transmitted by >40 whitefly species of the *B. tabaci* complex, the specifc combinations of virus-vector are numerous, how widespread is this mechanism involved in virion movement across midgut wall is yet to be discovered.

## Materials and Methods

### Insects, viruses and plants

Three cryptic species of the *B. tabaci* whitefly complex, Middle East Asia Minor 1 (MEAM1) (mitochondrial cytochrome oxidase I GenBank accession no. GQ332577.1), Mediterranean (MED) (mitochondrial cytochrome oxidase I GenBank accession no. GQ371165) and Asia II 1 (mitochondrial cytochrome oxidase I GenBank accession no. DQ309077) were reared on cotton plants (*Gossypium hirsutum* L. cv. Zhemian 1793) in insect-proof cages at 26 °C (± 1 °C) under a photoperiod of 14:10 h (light/dark) and relative humidity of 50% (± 10%). The purity of the culture was monitored every three generations by amplifying and sequencing the mitochondrial cytochrome oxidase I gene, which has been used widely to differentiate *B. tabaci* genetic groups (17). Clones of TYLCV isolate SH2 (GenBank accession no. AM282874.1) were agroinoculated into plants of tomato (*Solanum lycopersicum* L. cv. Hezuo903). Plants were grown in insect-proof greenhouses under controlled temperature at 25 ± 3 °C and natural lighting supplemented with artificial lights for 14 h a day from 0600 to 2000 hours.

### Tissue collection

For midgut and fat body isolation, the whiteflies were anesthetized on ice for 5 min, and then dissected from the abdomen in pre-chilled PBS buffer. The midguts and fat bodies were collected separately without contamination from other tissues and placed in PBS buffer. The whiteflies were then dissected from the prothorax for PSGs isolation. The midguts and PSGs were washed twice in PBS to remove contaminating viruses or proteins from the hemolymph. For hemolymph collection, each whitefly was dissected from the abdomen in 10 μL pre-chilled PBS buffer to release the content. Then all the liquid was collected without contamination from other tissues.

### Preparation of midgut protein and IP-UPLC-MS/MS

About 6,000 of newly emerged whiteflies were allowed to feed on TYLCV-infected tomato plants for one week, then midguts were dissected from these whiteflies and washed twice in PBS before collection. Total protein was extracted from 2,000 midguts using the lysis buffer supplied in the Capturem IP & Co-IP kit (Takara, 635721). The extracted proteins were divided into two equal parts and used for immunoprecipitation (IP), one with a TYLCV coat protein (CP) specific mouse monoclonal antibody and the other with a mouse pre-immune sera (control sera), using the Capturem IP & Co-IP kit according to the manufacturer’s instruction. The immunoprecipitate was then digested according to a filter aided sample preparation (FASP) method (45). The shotgun ultra-performance liquid chromatography-tandem MS (UPLC-MS/MS) procedure and data analysis were performed as previously reported (46). The MS/MS spectra were searched against the peptide database of MEAM1 species of *B. tabaci* (http://www.whiteflygenomics.org) using SEQUEST HT search engine configured with a Proteome Discoverer 1.4 workflow (Thermo Fischer Scientific, Bremen, Germany). The search parameters include 10 ppm and 0.8 Da mass tolerances for MS and MS/MS respectively, trypsin as the proteolytic enzyme with two allowed missed cleavage, oxidation and deamidated as dynamic modifications, and carbamidomethyl as static modification. Further, the peptides were extracted using high peptide confidence. 1% FDR (False discovery rate) was calculated using a decoy database by searching the peptide sequence.

### q(RT)-PCR analysis

Groups of 100 midguts or the fat bodies of 100 whiteflies were used to measure Vg expression levels in these tissues, and groups of 20 female or male whiteflies were used for Vg expression level determination in whitefly whole bodies. Total RNA was isolated using TRIzol reagent (Ambion, 15596018), and then cDNAs were produced using the PrimeScript RT reagent kit with gDNA Eraser (TaKaRa, RR047A). For viral DNA load quantification in whitefly whole bodies, total DNA was extracted from groups of 20 female or male whiteflies using previously described methods (47). Whitefly whole bodies were ground in 40 μL of ice-cold lysis buffer (50 mM Tris-HCl pH 8.4, 0.45% Tween 20, 0.45% Nonidet P-40, 0.2% gelatin and 60 mg/L proteinase K) and were incubated at 65 °C for 2 h, and then at 100 °C for 10 min. The supernatants were kept at -20 °C. For viral DNA load quantification in whitefly tissues, one PSG or the hemolymph of one female whitefly was dissected and ground in 10 μL of ice-cold lysis buffer, and then incubated at 65 °C for 4 h, and finally at 100 °C for 10 min. The supernatants were kept at -20 °C. q(RT)-PCR was performed using an ABI Prism 7500 Fast Real-Time PCR system (Applied Biosystems) with SYBR Premix Ex TaqTM II (TaKaRa, RR820A) and the primers are shown in S3 Table. For each reaction, 0.8 μL of each primer (10 mM), 6.4 μL of nuclease-free water, and 10 μL of SYBR Premix Ex Taq were added, in a total volume of 20 μL. The q(RT)-PCR protocol was 95 °C for 30 s, followed by 40 cycles of 95 °C for 5 s and 60 °C for 30 s. A negative control (nuclease-free water) was included throughout the experiments to detect contamination and to determine the degree of dimer formation. The results (threshold cycle values) of the q(RT)- PCR assays were normalized to the expression level of *B. tabaci β-actin* gene. The relative gene expression level or relative abundance of viral DNA was calculated using the 2^−Δ Ct^ method.

### Immunofluorescence assay

Whitefly tissues were dissected freshly and fixed in 4% paraformaldehyde (MultiSciences Biotech, LK-F0001) for 1 h at room temperature, and washed in TBST (TBS buffer with 0.05 % Tween 20) three times. The specimens were then permeabilized using 0.1% Triton X-100 in TBS for 1 h and blocked using TBST/BSA (TBST with 1% BSA) for 2 h at room temperature, followed by incubation with primary antibody in TBST/BSA overnight at 4 °C. The specimens were subsequently incubated with Dylight 488 (MultiSciences Biotech, LK-GAM4882) or Dylight 549 (MultiSciences Biotech, LK-GAR5492) labelled secondary antibody (1:500) in TBST/BSA for 1 h at room temperature after extensive washing with TBST. For intracellular vesicle or actin-based microvillar membrane and visceral muscle staining, the midguts were further incubated with Alex 647 lectin HPA (Invitrogen, L32454) or Dylight 647 phalloidin (Yeasen, 40762ES75) for 1 h at room temperature. The mouse anti-Vg monoclonal antibody (27) was provided by Prof. Gong-Ying Ye, and the mouse anti-TYLCV CP monoclonal antibody (48) and rabbit anti-TYLCV CP polyclonal antibody were provided by Prof. Jian-Xiang Wu, Zhejiang University. The other antibodies were rabbit anti-GFP monoclonal antibody (Abcam, ab183734) and rabbit anti-Rab5 polyclonal antibody (26). For confocal imaging, samples were mounted in fluoroshield mounting medium with DAPI (Abcam, ab104139) and imaged on a Zeiss LSM 780 confocal microscope (Zeiss, Germany). ImageJ was used to create the fluorescence intensity profiles with default parameters.

### Production of recombinant proteins in *Drosophila* cells

Sequences encoding VitN, DUF and vWD domains of MEAM1 whitefly Vg (GenBank accession no. GU332720) were amplified by reverse transcription PCR from adult whiteflies respectively. The primers are shown in S3 Table. The PCR products were subcloned into the pAc5.1/V5-His A vector (Invitrogen, V4110-20) with a GFP tag sequence at its 3’ end terminus. All plasmids were sequenced and transfected individually into *Drosophila Schneider* 2 (S2) cells using Lipofectamine 3000 Transfection reagent kit (Invitrogen, L3000008) according to the manufacturer’s instruction. The successful expression of these recombinant proteins was confirmed by appearance of autofluorescence of GFP (green) when visualized under a confocal microscope. Cells transfected with mock vector or vector only cloned with the GFP sequence were served as controls. 72 h after transfection with the relevant vectors, cells were harvested, pelleted at 500 g for 5 min and then washed twice with PBS. Proteins were extracted using Minute total protein extraction kit (Invent, SN-002) according to the manufacturer’s instruction.

### Recombinant protein/midgut *in vivo* binding assays

Midgut binding assays were performed as described previously (28). Briefly, adult whiteflies 2 days post-eclosion were fed with a solution containing individual recombinant protein mixed with buffer (PBS) (10% glycerol, 0.01% Chicago Sky Blue and 5mg / mL of BSA) for 4 h through a membrane feeding chamber. Then whiteflies were transferred to another feeding chamber containing a 15% sucrose solution for 6 h to remove unbound proteins in the midgut. Midguts were then dissected from female whiteflies for recombinant protein detection by immunofluorescence staining with a rabbit anti-GFP monoclonal antibody. Whiteflies fed with GFP were served as control.

### GST pull-down and western blotting assay

The fragment of TYLCV CP was amplified and cloned into pGEX-6p-1 for fusion with GST. Primers are listed in S3 Table. The recombinant protein was expressed in *Escherichia coli* strain BL21 and purified. The GST-CP was bound to glutathione Sepharose beads (GE Healthcare, 17- 5132-01) for 3 h at 4 °C, and the mixtures were centrifuged for 5 min at 100 × g, and the supernatants were discarded. For mapping of Vg domains that interact with TYLCV CP, the protein extracts of *Drosophila* S2 cells expressing the three recombinant Vg domains were added to the beads respectively and incubated for 2 h at 4 °C. After being centrifuged and washed five times with PBS, the beads-bound proteins were eluted by boiling in PAGE buffer for 5 min, and then the proteins were separated by 12% SDS/PAGE and detected by anti-GFP antibody.

To test the impact of anti-Vg antibody to the interaction between MEAM1 endogenous Vg and TYLCV CP, the non-viruliferous whitefly soluble protein extracts were prepared in cell lysis buffer (20 mM Tris·HCl, pH 7.5, 150 mM NaCl, 1% Triton X-100, 20 Mm *β*-glycerophosphate, 10 mM NaF, 1 mM PMSF, 1 mM sodium orthovanadate, 10 mg/mL leupeptin, 2 mg/mL aprotinin, 1 mM EDTA). Mouse anti-Vg monoclonal antibody and the corresponding mouse pre-immune sera (control sera) (Beyotime, A7028) as controls were incubated with the whitefly soluble protein extracts for 4 h at 4 °C and then the protein extracts were added to the beads and incubated for 2 h at 4 °C. To examine the interaction between MED and Asia II 1 endogenous Vg and TYLCV CP, the soluble protein extracts form these two species were added to the beads and incubated for 2 h at 4 °C. The beads-bound proteins were eluted and detected by anti-Vg antibody as described above.

For protein detection in tissues or whole bodies of whiteflies, total protein was isolated from groups of 200 midguts, fat bodies of 200 female whiteflies or 100 whole whiteflies using the cell lysis buffer. Protein samples were separated by 12% SDS/PAGE, and transferred to polyvinylidine difluoride membranes. The membranes were blocked with 5% nonfat milk in phosphate-buffered saline (PBS; Sangon Biotech, SB0627) with 0.1% Tween 20 (BBI Life Sciences, 9005-64-5) and then incubated with the anti-TYLCV CP, anti-Vg or anti-actin (EarthOx, E021020-02) antibodies. After incubation with secondary antibody (MultiSciences Biotech, GAM007), signals were visualized with the ECL Plus Detection System (Bio-Rad, 170-5060).

### dsRNA preparation

dsRNA specific to Vg of MEAM1 (GenBank accession no. GU332720.1), MED (GenBank accession no. GU332722.1) or Asia II 1 (GenBank accession no. GU332721.1) was synthesized using the AmpliScribe T7- Flash Transcription Kit (Epicentre, ASF3507), following the manufacturer’s instructions. Briefly, the DNA template for dsRNA synthesis was amplified with primers containing the T7 RNA polymerase promoter at both ends (S3 Table), and the purified DNA template was then used to generate dsRNAs. dsRNA specific to GFP was synthesized as control. Subsequently, the synthesized dsRNA was purified via phenol-chloroform precipitation and resuspended in nuclease-free water, and the concentration of dsRNA was quantified with a NanoDrop 2000 (Thermo Fisher Scientific). Finally, the quality and size of the dsRNAs were further verified via electrophoresis in a 2% agarose gel.

### Gene silencing via oral ingestion of dsRNA

RNA silencing was performed as previously described (49). Briefly, dsRNAs were diluted into 15% (wt/vol) sucrose solution at the concentration of 300 ng/μL. Approximately 100 adult whiteflies at 1- 2 DAE were released into each feeding chamber. The tube was incubated in an insect-rearing room for 48 h. Subsequently, RNA was extracted from 20 female or male individuals to examine the gene expression level, and total proteins were extracted from 30 female individuals to examine the Vg protein level. The remaining insects were transferred to virus-infected plants for virus acquisition efficiency test. To compare the virus acquisition efficiency, the dsVg or dsGFP-treated whiteflies were placed to feed on two opposite leaflets of a TYLCV-infected tomato plant for 24 h, using leaf clip cages (50) (Fig. S7A). Then, the whiteflies were collected and used for quantitative assays, immunostaining assays and virus transmission tests. Each set of experiment was repeated three times.

### Oral ingestion of anti-Vg antibody

Adult whiteflies at 3-4 DAE were collected and fed with anti-Vg antibodies (1:100) or mouse pre- immune serum (1:100, control) in 15% (wt/vol) sucrose solution using membrane feeding device. Approximately 100 adult whiteflies were released into each feeding chamber, and the feeding device was incubated in an insect-rearing room for 48 h. Subsequently, various tissues were dissected from female whiteflies for Vg antibody detection. The remaining insects were transferred to virus-infected plants for virus acquisition efficiency test. To examine the virus infection rate of whiteflies during continuous feeding, insects were collected at 1, 2, 4, 6, 12 and 24 h after transfer and total DNA was extracted from single females and then subjected to PCR analysis according to previous descriptions (24, 51). For each time point, 24 female whiteflies were collected in total, and three replicates were conducted. In addition, non-viruliferous whiteflies were used as controls. To determine the virus abundance in whitefly whole body and various tissues, insects were collected after a 24 h AAP on virus-infected plants and used for quantitative assays, immunostaining assays and virus transmission tests. Each set of experiment was repeated three times.

### Transmission of TYLCV to plants by whiteflies

In the TYLCV transmission tests, whiteflies after dsRNAs feeding were given a 24 h AAP on TYLCV-infected plants and then the whiteflies were inoculated singly to individual uninfected tomato plants. The inoculation was performed on the top second leaf of the plant at the 3-4 true-leaf stage (∼3 weeks after sowing) for a 72 h inoculation access period (IAP), using a leaf clip cage (50). For TYLCV transmission by anti-Vg antibody treated whiteflies, viruliferous female whiteflies were inoculated singly to individual uninfected tomato plants as described above. For infectivity comparison of female and male adults at different developmental stages following a 48 h AAP, two females or males were used to inoculate one uninfected plant as described above. The plants were then sprayed with imidacloprid at a concentration of 20 mg/L to kill all the whitefly adults and eggs, and maintained in insect-proof cages at 26 °C (± 1 °C) under a photoperiod of 14:10 h (light/dark) to allow observation of disease symptoms. Thirty uninfected tomato plants were used for each transmission test.

### Statistical analysis

Data were presented as mean ± SEM of three independent biological replicates, unless otherwise noted. All analyses were performed using SPSS (version 13) software. Differences between the virus quantity in the hemolymph and PSGs of whiteflies were analyzed by the non-parametric Mann- Whitney U test. The others were assessed with independent-samples t-test.

## Acknowledgements

Financial support for this study was provided by the National Natural Science Foundation of China (31925033, 31930092). We thank Prof. Gong-Ying Ye for providing anti-Vg antibody and Prof. Jian- Xiang Wu for providing anti-TYLCV antibody.

## References

1. Sappington TW, Raikhel AS. Molecular characteristics of insect vitellogenins and vitellogenin receptors. Insect Biochem Mol Biol. 1998;28:277–300.

2. Snigirevskaya ES, Raikhel AS. Receptor-mediated endocytosis of yolk proteins in insect oocytes. In: Raikhel AS, editor. Reproductive biology of invertebrates: Progress in vitellogenesis. Enfield, NH: Science Publishers, Inc.; 2005. p. 199–228.

3. Chen JS, Sappington TW, Raikhel AS. Extensive sequence conservation among insect, nematode, and vertebrate vitellogenins reveals ancient common ancestry. J Mol Evol. 1997;44:440–51.

4. Tufail M, Takeda M. Molecular characteristics of insect vitellogenins. J Insect Physiol. 2008;54:1447–58.

5. Giorgi F, Snigirevskaya E, Raikhel AS. The cell biology of yolk protein precursor synthesis and secretion. In: Raikheil AS, editor. Reproductive biology of invertebrates: Progress in vitellogenesis. Enfield, NH: Science Publisher Inc.; 2005. p. 33–68.

6. Seehuus SC, Norberg K, Krekling T, Fondrk K, Amdam GV. Immunogold localization of vitellogenin in the ovaries, hypopharyngeal glands and head fat bodies of honeybee workers, *Apis mellifera*. J Insect Sci. 2007;7:1–14.

7. Li JL, Huang JX, Cai WZ, Zhao ZW, Peng WJ, Wu J. The vitellogenin of the bumblebee, *Bombus hypocrita*: studies on structural analysis of the cDNA and expression of the mRNA. J Comp Physiol B. 2010;180:161–70.

8. Huo Y, Yu YL, Chen LY, Li Q, Zhang MT, Song ZY, et al. Insect tissue-specific vitellogenin facilitates transmission of plant virus. PLoS Pathog. 2018;14:e1006909.

9. Gray SM, Banerjee N. Mechanisms of arthropod transmission of plant and animal viruses. Microbiol Mol Biol Rev. 1999;63:128–48.

10. Hogenhout SA, Ammar ED, Whitfield AE, Redinbaugh MG. Insect vector interactions with persistently transmitted viruses. Annu Rev Phytopathol. 2008;46:327–59.

11. Nagata T, Inoue-Nagata AK, van Lent J, Goldbach R, Peters D. Factors determining vector competence and specificity for transmission of *Tomato spotted wilt virus*. J Gen Virol. 2002;83:663–71.

12. Ammar ED, Gomez-Luengo RG, Gordon DT, Hogenhout SA. Characterization of *Maize iranian mosaic virus* and comparison with Hawaiian and other isolates of *Maize mosaic virus* (Rhabdoviridae). J Phytopathol. 2005;153:129–36.

13. Merrill MH, Tenbroeck C. The transmission of equine encephalomyelitis virus by *Aedes aegypti*. J Exp Med. 1935;62:687–95.

14. Storey HH. Investigations of the mechanism of the transmission of plant viruses by insect vectors. Proc Biol Sci. 1933;113:463–85.

15. Blanc S, Drucker M, Uzest M. Localizing viruses in their insect vectors. Annu Rev Phytopathol. 2014;52:403–25.

16. Neelakanta G, Sultana H. Viral receptors of the gut: vector-borne viruses of medical importance. Curr Opin Insect Sci. 2016;16:44–50.

17. De Barro PJ, Liu SS, Boykin LM, Dinsdale AB. *Bemisia tabaci*: A statement of species status. Annu Rev Entomol. 2011;56:1–19.

18. Rosen R, Kanakala S, Kliot A, Pakkianathan BC, Abu Farich B, Santana-Magal N, et al. Persistent, circulative transmission of begomoviruses by whitefly vectors. Curr Opin Virol. 2015;15:1–8.

19. Navas-Castillo J, Fiallo-Olivé E, Sánchez-Campos S. Emerging virus diseases transmitted by whiteflies. Annu Rev Phytopathol 2011;49:219–48.

20. Rojas MR, Macedo MA, Maliano MR, Soto-Aguilar M, Souza JO, Briddon RW, et al. World management of geminiviruses. Annu Rev Phytopathol. 2018;56:637–77.

21. Czosnek H. Tomato yellow leaf curl virus disease: management, molecular biology, breeding for resistance. Czosnek H, editor. Netherlands: Springer 2007.

22. Scholthof KBG, Adkins S, Czosnek H, Palukaitis P, Jacquot E, Hohn T, et al. Top 10 plant viruses in molecular plant pathology. Mol Plant Pathol. 2011;12:938–54.

23. Ghanim M. A review of the mechanisms and components that determine the transmission efficiency of *Tomato yellow leaf curl virus (Geminiviridae; Begomovirus*) by its whitefly vector. Virus Res. 2014;186:47–54.

24. Wei J, Zhao JJ, Zhang T, Li FF, Ghanim M, Zhou XP, et al. Specific cells in the primary salivary glands of the whitefly *Bemisia tabaci* control retention and transmission of begomoviruses. J Virol. 2014;88:13460–8.

25. Pan LL, Chen QF, Zhao JJ, Guo T, Wang XW, Hariton-Shalev A, et al. Clathrin-mediated endocytosis is involved in *Tomato yellow leaf curl virus* transport across the midgut barrier of its whitefly vector. Virology. 2017;502:152–9.

26. Xia WQ, Liang Y, Chi Y, Pan LL, Zhao J, Liu SS, et al. Intracellular trafficking of begomoviruses in the midgut cells of their insect vector. PLoS Pathog. 2018;14:e1006866.

27. Guo JY, Dong SZ, Yang XL, Cheng L, Wan FH, Liu SS, et al. Enhanced vitellogenesis in a whitefly via feeding on a begomovirus-infected plant. PLoS One. 2012;7:e43567.

28. Wang LL, Wei XM, Ye XD, Xu HX, Zhou XP, Liu SS, et al. Expression and functional characterisation of a soluble form of tomato yellow leaf curl virus coat protein. Pest Manag Sci. 2014;70:1624–31.

29. Ghanim M, Rosell RC, Campbell LR, Czosnek H, Brown JK, Ullman DE. Digestive, salivary, and reproductive organs of *Bemisia tabaci* (Gennadius) (Hemiptera: Aleyrodidae) B type. J Morphol. 2001;248:22–40.

30. Sanchez JF, Lescar J, Chazalet V, Audfray A, Gagnon J, Alvarez R, et al. Biochemical and structural analysis of *Helix pomatia* agglutinin: A hexameric lectin with a novel fold. J Biol Chem. 2006;281:20171–80.

31. Don-Wheeler G, Engelmann F. The biosynthesis and processing of vitellogenin in the fat bodies of females and males of the cockroach *Leucophaea maderae*. Insect Biochem Mol Biol. 1997;27:901–18.

32. Royzokan EM, Cunningham CB, Hebb LE, Mckinney EC, Moore AJ. Vitellogenin and vitellogenin receptor gene expression is associated with male and female parenting in a subsocial insect. Proc Biol Sci. 2015;282:20150787.

33. Martinez T, Wheeler D. Identification of vitellogenin in the ant, *Camponotus festinatus*: changes in hemolymph proteins and fat body development in workers. Arch Insect Biochem Physiol. 1991;17:143–55.

34. Ning WX, Shi Xb, Liu Bm, Pan Hp, Wei Wt, Zeng Y, et al. Transmission of *Tomato yellow leaf curl virus* by *Bemisia tabaci* as affected by whitefly sex and biotype. Sci Rep. 2015;5:10744.

35. Rom M, Antignus Y, Gidoni D, Pilowsky M, Cohen S. Accumulation of tomato yellow leaf curl virus DNA in tolerant and susceptible tomato lines. Plant Dis. 1993;77:253–7.

36. Dow JAT. Insect midgut function. Adv In Insect Phys. 1986;19:189–328.

37. Roth Z, Weil S, Aflalo ED, Manor R, Sagi A, Khalaila I. Identification of receptor-interacting regions of vitellogenin within evolutionarily conserved beta-sheet structures by using a peptide array. Chembiochem. 2013;14:1116–22.

38. Li AK, Sadasivam M, Ding JL. Receptor-ligand interaction between vitellogenin receptor (VtgR) and vitellogenin (Vtg), implications on low density lipoprotein receptor and apolipoprotein B/E. J Biol Chem. 2003;278:2799–806.

39. Tufail M, Raikhel AS, Takeda M. Biosynthesis and processing of insect vitellogenins. In: Raikheil AS, editor. Reproductive biology of invertebrates: Progress in vitellogenesis. 12. Enfield, NH: Science Publishers, Inc.; 2005. p. 1–32.

40. Pennisi E. Armed and dangerous. Science. 2010;327:804–5.

41. Chen Q, Wei TY. Viral receptors of the gut: insect-borne propagative plant viruses of agricultural importance. Curr Opin Insect Sci. 2016;16:9–13.

42. Fu H, Leake CJ, Mertens PPC, Mellor PS. The barriers to bluetongue virus infection, dissemination and transmission in the vector, *Culicoides variipennis* (Diptera : Ceratopogonidae). Arch Virol. 1999;144:747–61.

43. Grove J, Marsh M. The cell biology of receptor-mediated virus entry. J Cell Biol. 2011;195:1071–82.

44. Tufail M, Takeda M. Insect vitellogenin/lipophorin receptors: molecular structures, role in oogenesis, and regulatory mechanisms. J Insect Physiol. 2009;55:88–104.

45. Wiśniewski JR. Quantitative evaluation of filter aided sample preparation (FASP) and multienzyme digestion FASP protocols. Anal Chem. 2016;88:5438–43.

46. Huang HJ, Lu JB, Li Q, Bao YY, Zhang CX. Combined transcriptomic/proteomic analysis of salivary gland and secreted saliva in three planthopper species. J Proteomics. 2018;172:25–35.

47. Luo C, Yao Y, Wang RJ, Yan FM, Hu DX, Zhang ZL. The use of mitochondrial cytochrome oxidase I (mt CO I) gene sequences for the identification of biotypes of *Bemisia tabaci* (Gennadius) in China. Acta Entomol Sinica. 2002;45:757–63.

48. Wu JX, Shang HL, Xie Y, Shen QT, Zhou XP. Monoclonal antibodies against the whitefly-transmitted tomato yellow leaf curl virus and their application in virus detection. J Integr Agric. 2012;11:263–8.

49. Wei J, He YZ, Guo Q, Guo T, Liu YQ, Zhou XP, et al. Vector development and vitellogenin determine the transovarial transmission of begomoviruses. Proc Natl Acad Sci U S A. 2017;114:6746–51.

50. Jiu M, Zhou XP, Liu SS. Acquisition and transmission of two begomoviruses by the B and a non-B biotype of *Bemisia tabaci* from Zhejiang, China. J Phytopathol. 2010;154:587–91.

51. Ghanim M, Morin S, Czosnek H. Rate of *Tomato yellow leaf curl virus* translocation in the circulative transmission pathway of its vector, the whitefly *Bemisia tabaci*. Phytopathology. 2001;91:188–96.

