## Supplemental Figures and Tables for "Gut expressed vitellogenin is hijacked by a virus to facilitate its spread"

**Supplementary materials**

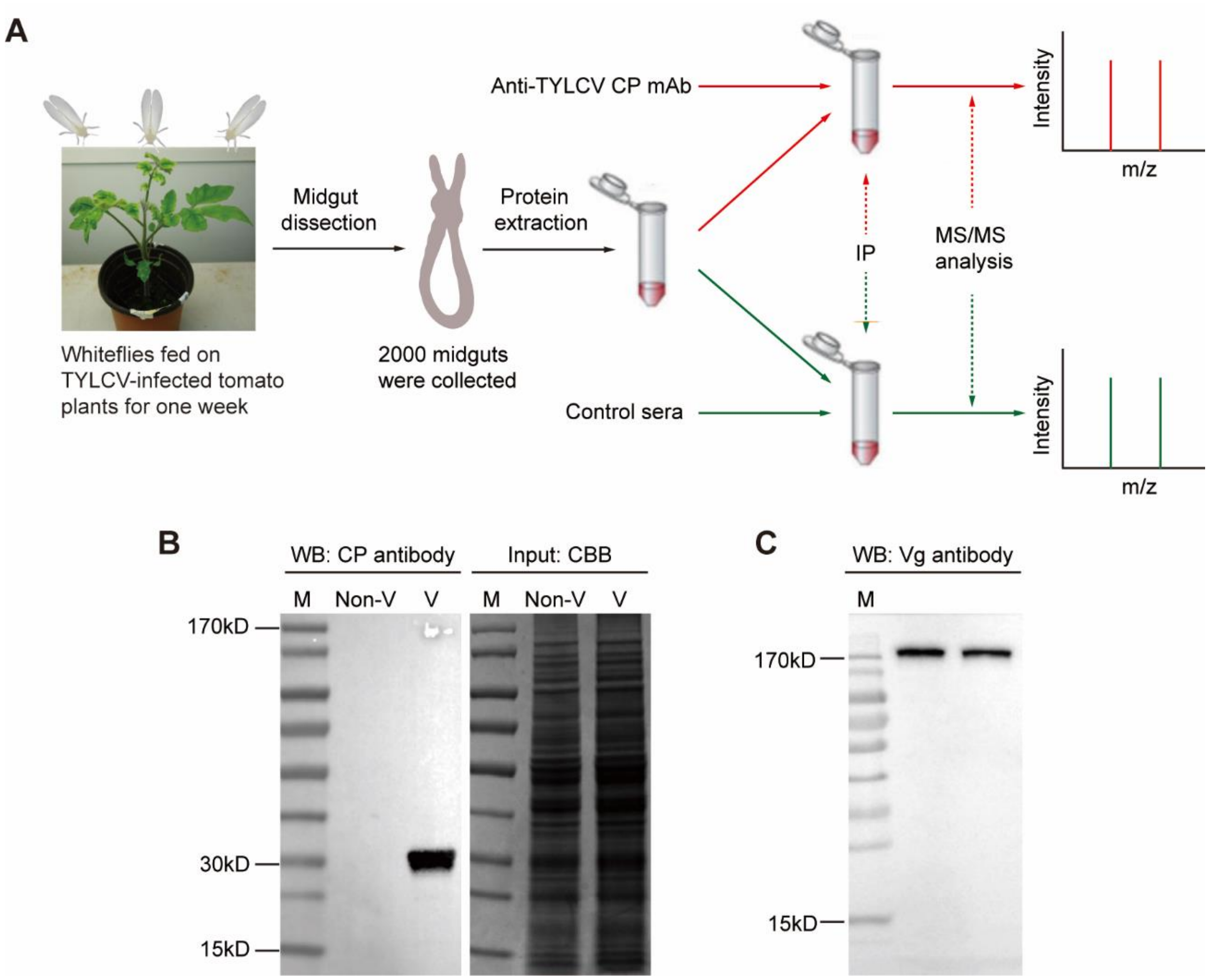


**Fig S1. Schematic representation of the study design (A) and antibody specificity test.** About 6000 newly emerged whiteflies were allowed to feed on TYLCV-infected tomato plants for one week, then, 2000 midguts were dissected from these whiteflies and used for protein extraction. The extracted proteins were divided into two equal parts and used for immunoprecipitation (IP), one with a mouse anti-TYLCV coat protein (CP) monoclonal antibody (mAb) and the other with mouse pre-immune sera (control sera). The immunoprecipitate was further digested with trypsin and analyzed by shotgun ultra-performance liquid chromatography (UPLC)-tandem MS (MS/MS). (B) Anti-TYLCV CP monoclonal antibody specificity test. A single band was detected by the anti-TYLCV CP antibody in total protein extract from viruliferous (V) but not in that from nonviruliferous (Non-V) whiteflies. Coomassie Brilliant Blue (CBB) staining of total proteins serves as a loading control. (C) Anti-Vg monoclonal antibody specificity test. A single band was detected by the anti-Vg monoclonal antibody in total protein extract from adult whiteflies. M, marker.

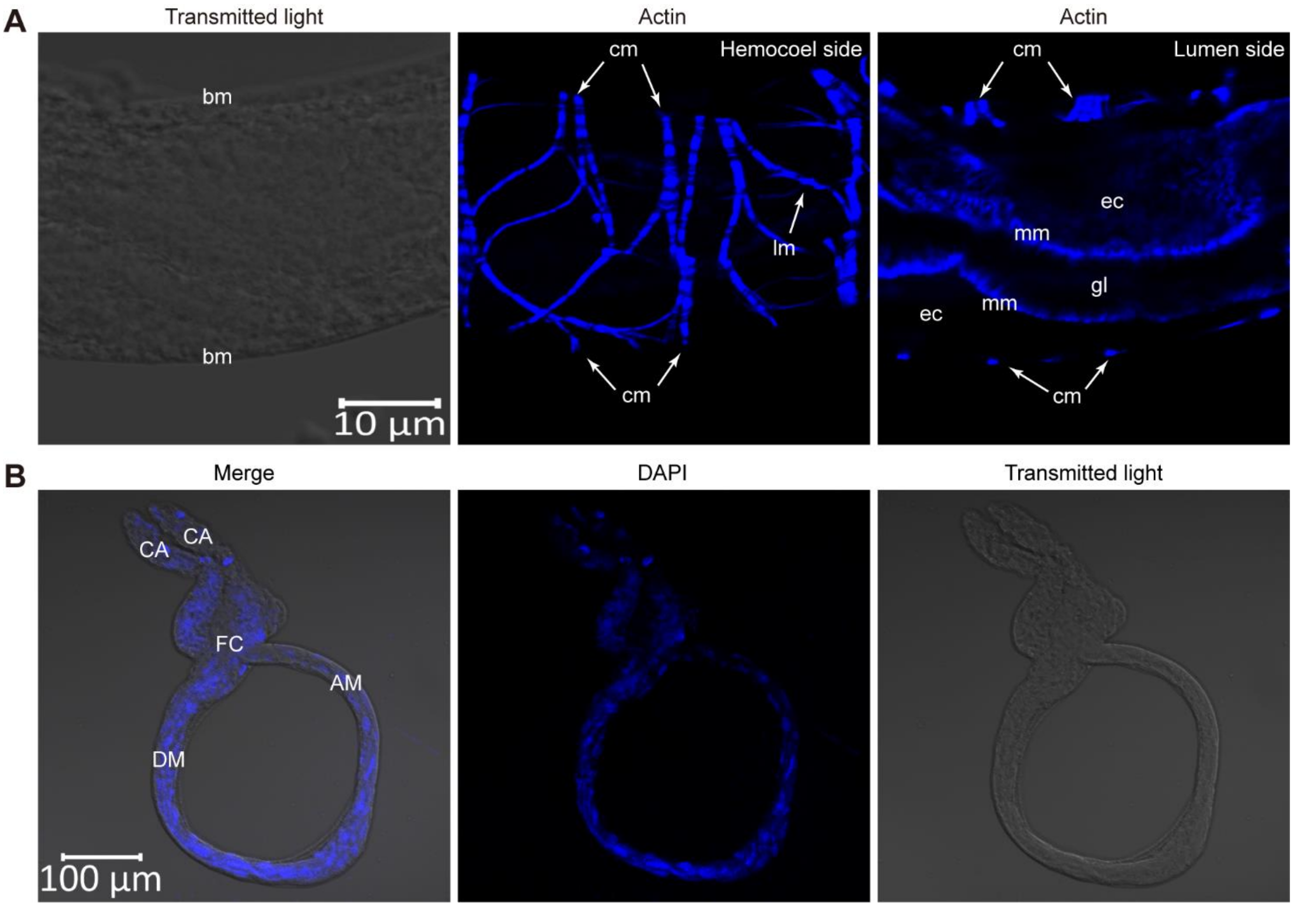


**Fig S2. Structure of whitefly midgut.** (A) Alimentary canal structure of whitefly, composing of a single layer of epithelial cells, with microvillar membrane on the lumen side and basal membrane on the hemocoel side, covered with circular and longitudinal muscles. The actin-based microvillar membrane and muscle fibers were stained with Dylight 647 phalloidin (blue). *bm* basal membrane, *cm* circular muscle, *lm* longitudinal muscle, *gl* gut lumen, *mm* microvillar membrane, *ec* epithelial cell. (B) Midgut of whitefly, comprising the gastric caecum (CA), filter chamber (FC), descending midgut (DM) and ascending midgut (AM). Cell nucleus was stained with DAPI (blue). Images are representative of multiple experiments with multiple preparations.

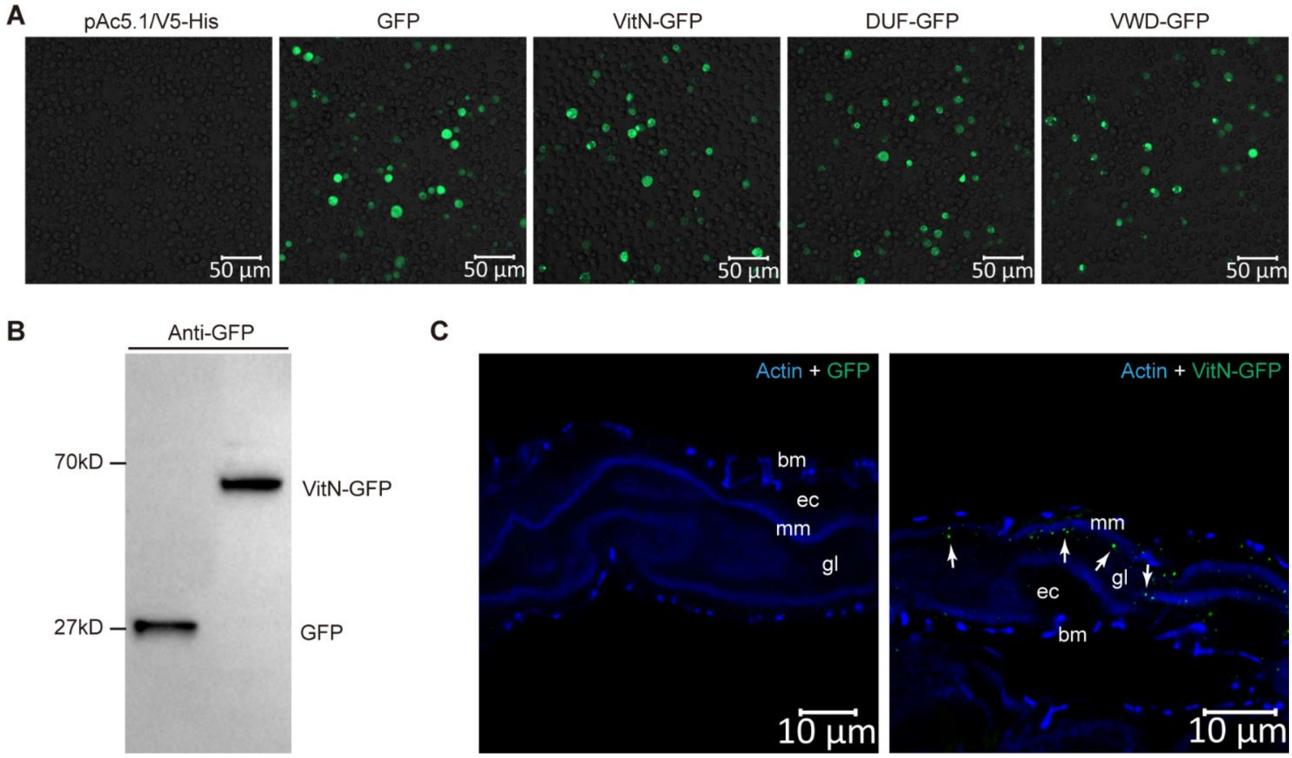


**Fig S3. Expression of Vg domains for in vivo midgut binding assays.** (A) Expression of GFP, VitN-GFP, DUF-GFP and vWD-GFP in Drosophila Schneider 2 (S2) cells. The pAc5.1/V5-His mock vector was used as control. The cells were visualized using a Zeiss LSM 780 confocal microscope with a 488-nm laser for GFP autofluorescence (green) excitation. (B) Western blotting of GFP and recombinant VitN-GFP using a rabbit anti-GFP monoclonal antibody. (C) The VitN-GFP is able to bind to whitefly midgut (right) but the GFP alone is unable to (left). Whiteflies were fed with GFP or VitN-GFP for 4 h followed by a 6 h feeding on a sucrose solution to remove unbound proteins. Then midguts were dissected for immunostaining using a rabbit anti-GFP monoclonal antibody and goat anti-rabbit IgG labeled with Dylight 488 (green) secondary antibody. The white arrow indicates the immune-reactive signal of the VitN-GFP protein. gl gut lumen, mm microvillar membrane, ec epithelial cell, bm basal membrane. Images are representative of three independent experiments with a total of 30 whiteflies analyzed for each treatment.

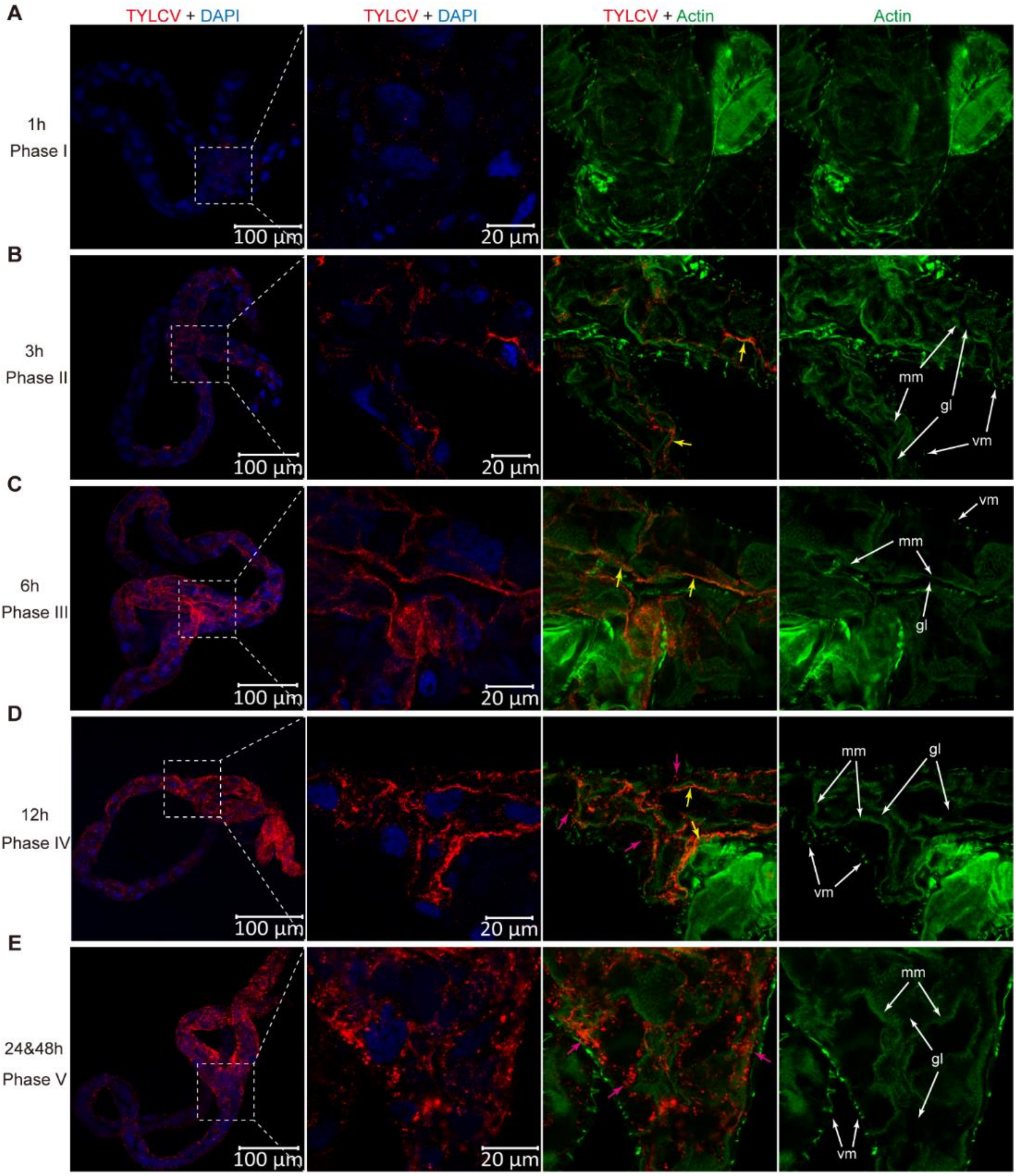


**Fig S4. Infection process of TYLCV in the midgut of whiteflies.** Midguts of female whiteflies feeding on TYLCV-infected tomato plants for 1, 3, 6, 12, 24 and 48 h AAP were dissected and prepared for immunofluorescence. After 1 h AAP, virus signals were only detected in the filter chamber (A, phase I), or also detected in the gastric caecum and descending midgut (B, phase II). After 3 h AAP, most of the midguts appeared as phase II, and after 6 h APP, TYLCV virions were seen throughout the midgut (C, phase III). In phase II and phase III, the virions were bound to the microvillar membrane and not seen in the cytoplasm. After 12 h AAP, some virions were seen in the cytoplasm of epithelial cells (D, phase IV). After 24 and 48 h AAP, virions were seen in the cytoplasm close to the basal membrane of epithelial cells (E, phase V). TYLCV was detected using a mouse anti-CP monoclonal antibody and goat anti-mouse IgG labeled with Dylight 549 (red) secondary antibody. Cell nucleus was stained with DAPI (blue). Actin-based microvillar membrane and visceral muscles were stained with Dylight 647 phalloidin (green). Yellow arrow indicates TYLCV virions bound to the microvillar membrane. Reddish-purple arrow indicates TYLCV virions that have invaded the cytoplasm. *gl* gut lumen, *mm* microvillar membrane, *vm* visceral muscles. Images are representative of a total of 30 midguts examined for each time point.

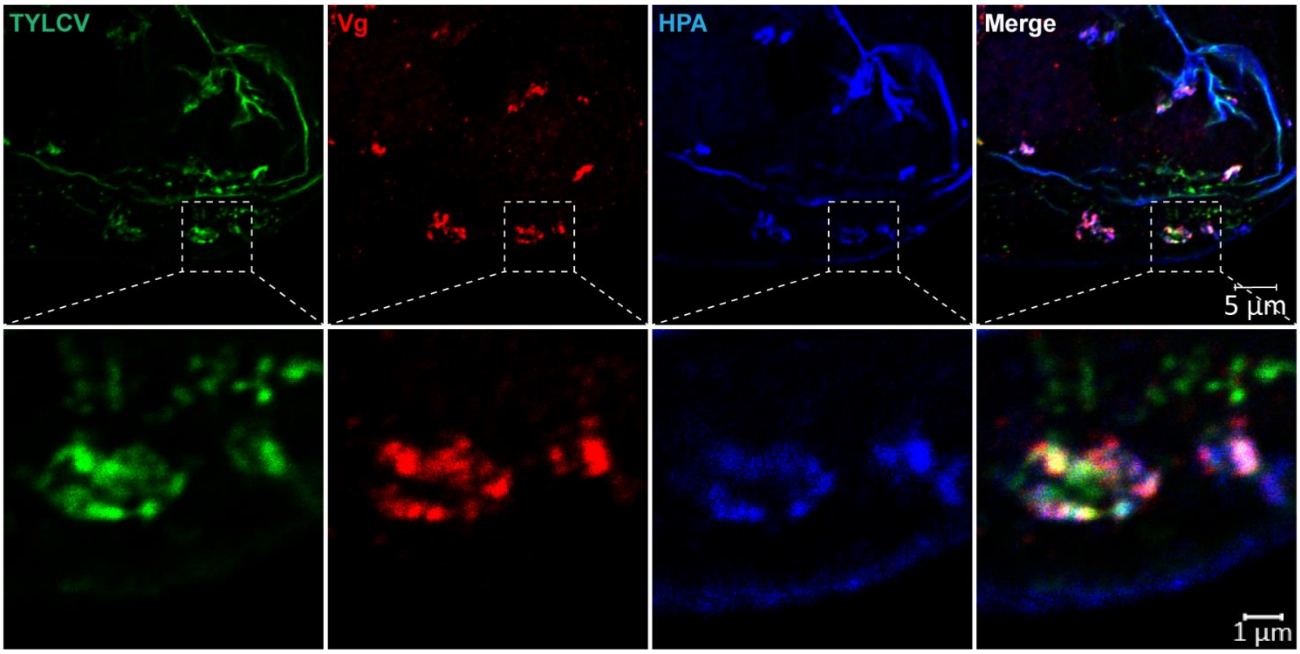


**Fig S5. Co-localization of TYLCV, Vg and lectin HPA-labelled vesicles in the midgut epithelial cells.** Midguts of female whiteflies exposed to TYLCV-infected plants for a 72 h AAP were dissected and used for immunofluorescence. TYLCV was detected using a rabbit anti-CP polyclonal antibody and goat anti-rabbit IgG labelled with Dylight 488 (green) secondary antibody. Vg was detected using a mouse anti-Vg monoclonal antibody and goat anti-mouse IgG labelled with Dylight 549 (green) secondary antibody. Intracellular vesicles were labelled using Alex 647 lectin HPA (blue). Images are representative of multiple experiments with multiple preparations.

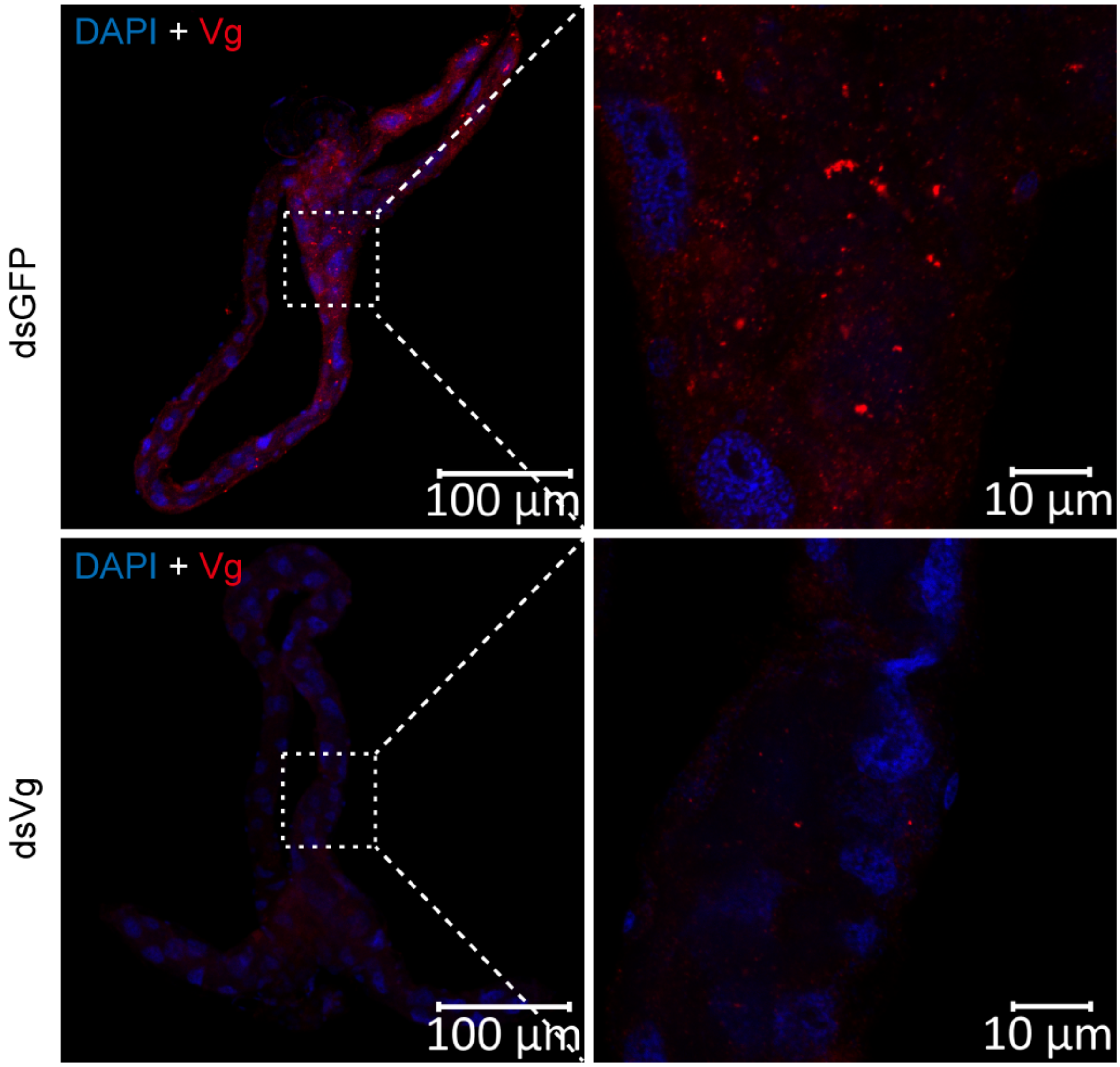


**Fig S6. Localization of Vg protein in the midguts of dsGFP or dsVg treated whiteflies.** Vg was detected using a mouse anti-Vg monoclonal antibody and goat anti-mouse IgG labeled with Dylight 549 (red) secondary antibody. Cell nucleus was stained with DAPI (blue). Images are representative of three independent experiments with a total of 20 whiteflies analyzed for each treatment.

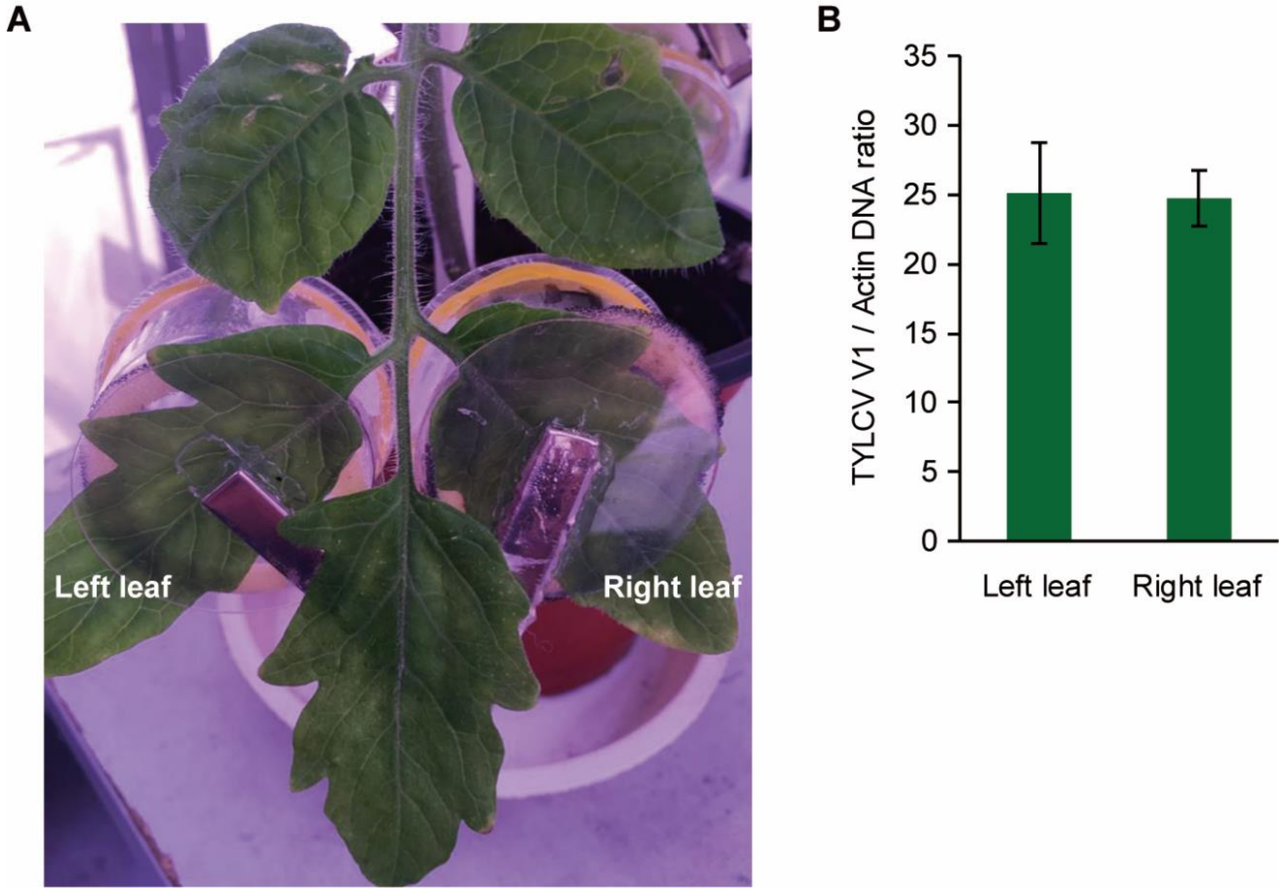


**Fig S7. Schematic representation of the method used to compare virus acquisition efficiency of whiteflies.** (A) Two groups of whiteflies were put on the opposite leaves of a TYLCV-infected tomato plant using leaf clip cages. (B) Similar virus amounts were acquired by whiteflies after a 24 h AAP on the two opposite leaves (left leaf and right leaf). Mean ± SEM from three independent experiments.

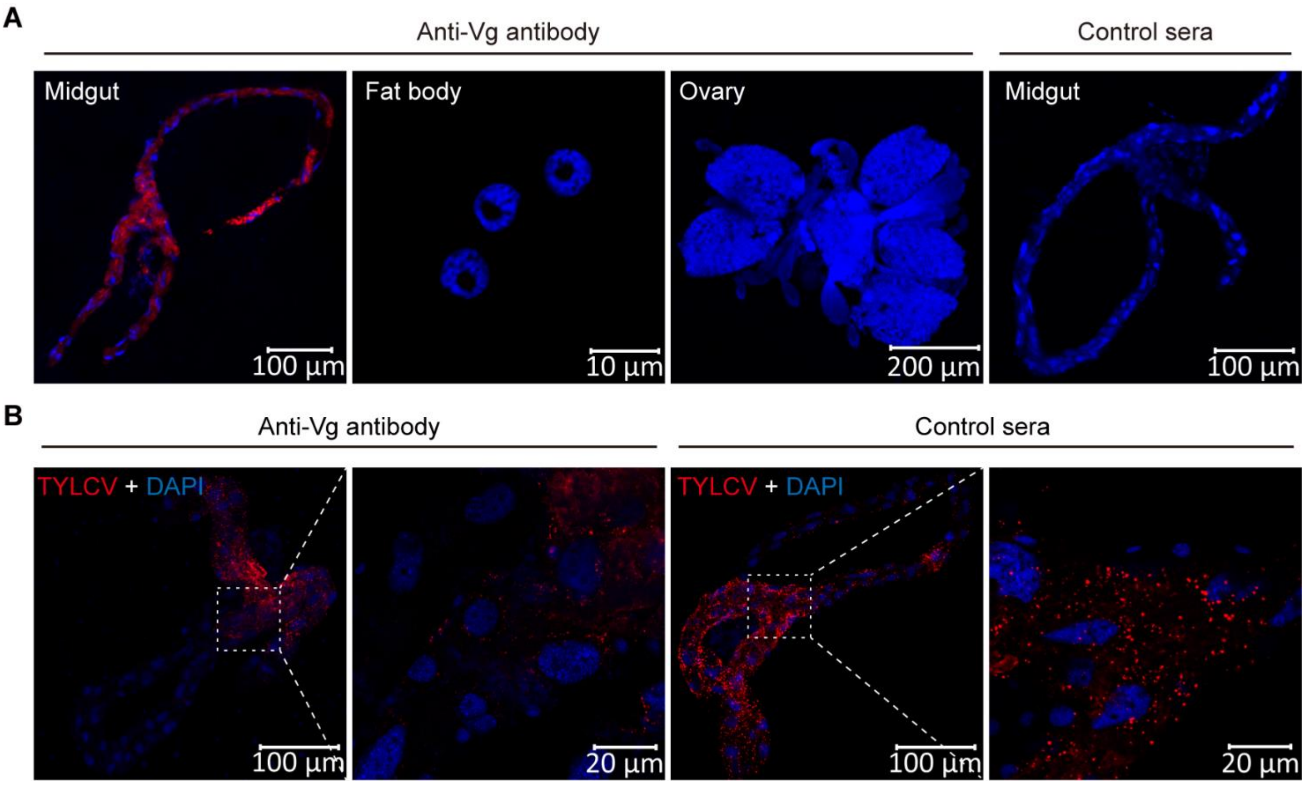


**Fig S8. Immune-blocking of midgut Vg inhibited the movement of TYLCV across the midgut wall.** (A) Detection of ingested antibody in various tissues of whiteflies. Adult female whiteflies were fed with mouse anti-Vg monoclonal antibody or mouse pre-immune serum (control serum) for 48 h via membrane feeding. Then midguts, fat bodies and ovaries were dissected and prepared for immunofluorescence. The Vg antibody and control serum were detected using a goat anti-mouse IgG labeled with Dylight 549 (red). Cell nucleus was stained with DAPI (blue). (B) Localization of TYLCV in midguts of anti-Vg antibody or control sera treated whiteflies after a 24 h AAP on TYLCV-infected tomato plants. TYLCV was detected using a mouse anti-CP monoclonal antibody and goat anti-mouse IgG labeled with Dylight 549 (red) secondary antibody. Cell nucleus was stained with DAPI (blue). Images are representative of three independent experiments with a total of 30 whiteflies analyzed for each experiment.

**Table S1.** Detection of TYLCV coat protein in the midguts of whiteflies by immunofluorescence microscopy at various times following the first access of whitefly to TYLCV-infected tomato plants.

| Hours post first access of whitefly to TYLCV-infected tomato | % Midguts of whitefly with virions in each of the five phases (n=30)* | | | | |
|---|---|---|---|---|---|
| | **Phase I** | **Phase II** | **Phase III** | **Phase IV** | **Phase V** |
| 0 | 0 | 0 | 0 | 0 | 0 |
| 1 | 23 | 40 | 7 | 0 | 0 |
| 3 | 7 | 60 | 17 | 7 | 0 |
| 6 | 3 | 40 | 44 | 13 | 0 |
| 12 | 0 | 17 | 13 | 60 | 10 |
| 24 | 0 | 3 | 7 | 40 | 50 |
| 48 | 0 | 0 | 0 | 7 | 93 |

*Phase I: TYLCV virions were only detected in the filter chamber of midgut;

Phase II: virions were present in the filter chamber, gastric caecum and descending midgut;

Phase III: virions bound to microvilli throughout the whole midgut;

Phase IV: some virions have moved to the cytoplasm of midgut epithelial cells;

Phase V: most of the virions have moved to the cytoplasm to spread into the hemolymph.

**Table S2.** Ingestion of dsVg or anti-Vg antibody via membrane feeding inhibited viral movement in the midgut and transmission by whiteflies.

| **Treatment** | | **% Midguts of whitefly with virions in each of the five phases following 24 h acquisition on TYLCV-infected tomato** | | | | | **% Tomato plants show disease symptoms at 15 and 30 days post transmission (n=30)** | |
|---|---|---|---|---|---|---|---|---|
| | ***N*.*** | **Phase I** | **Phase II** | **Phase III** | **Phase IV** | **Phase V** | **15 d** | **30 d** |
| dsGFP | 60 | 0 | 3 | 10 | 36 | 51 | 37 | 43 |
| dsVg | 60 | 0 | 7 | 22 | 45 | 26 | 17 | 30 |
| Control sera | 30 | 0 | 2 | 10 | 33 | 55 | 40 | 57 |
| Vg antibody | 30 | 0 | 6 | 24 | 40 | 30 | 13 | 22 |

*The total number of midguts examined for each treatment.

### Table S3. Primers used in this study

| Primer name | Sequence (5'-3')* | Length (bp) | Purpose |
|---|---|---|---|
| TYLCV-F | ATACCTGGACACCTAATGGC | 413 | TYLCV DNA detection |
| TYLCV-R | AGTCACGGGCCCTTACA | | |
| qTYLCV-F | GAAGCGACCAGGCGATATAA | 189 | q-PCR for TYLCV total DNA |
| qTYLCV-R | GGAACATCAGGGCTTCGATA | | |
| qMEAM1 Vg-F | ACAAGTCTCCGACGCCGAAG | 185 | qRT-PCR for MEAM1 *Vg* |
| qMEAM1 Vg -R | TTGACATCGGCTTTACGGCA | | |
| qMED Vg-F | TACGCTGACACTTCACCGCC | 183 | qRT-PCR for MED *Vg* |
| qMED Vg -R | GTCGCTGCGGCTAACGTAGT | | |
| qAsia II 1 Vg-F | CTGCCGTCTACGCTGTCGTT | 125 | qRT-PCR for Asia II 1 *Vg* |
| qAsia II 1 Vg -R | GTCGCTGCGGCTAACGTAGT | | |
| qβ-Actin-F | TCTTCCAGCCATCCTTCTTG | 173 | q(RT)-PCR for *β-Actin* |
| qβ-Actin-R | CGGTGATTTCCTTCTGCATT | | |
| MEAM1 Vg-RNAi-F | T7-ACATCGTCAAGGCCACCAA | 451 | MEAM1 *Vg* dsRNA synthesis |
| MEAM1 Vg-RNAi-R | T7-TAGAGCTGGAACTAGATGAG | | |
| MED Vg-RNAi-F | T7-TAGCAGCGACTCCAGCTCCT | 275 | MED *Vg* dsRNA synthesis |
| MED Vg-RNAi-R | T7-CGGGCTTGGCTGGGTATCTG | | |
| Asia II 1 Vg-RNAi-F | T7-TAGCAGCGACTCCAGCTCCT | 275 | Asia II 1 *Vg* dsRNA synthesis |
| Asia II 1 Vg-RNAi-R | T7-CGGGCTTGGCTGGGTATCTG | | |
| gfp-RNAi-F | T7-CTCGTGACCACCCTGACCTAC | 314 | *gfp* dsRNA synthesis |
| gfp-RNAi-R | T7-GTTCACCTTGATGCCGTTCTT | | |
| GFP-F | ATTTGCGGCCGCATGGTGAGCAAGGGCGAG | 720 | Expression of GFP in S2 cells |
| GFP-R | CCCTCGAGCTTGTACAGCTCGTCCATGC | | (*Not* I and *Xho* I sites are underlined) |
| VitN-GFP-F | CGGAATTCACTCCTTTCTTCCCCTAC | 1029 | Expression of VitN-GFP in S2 cells |
| VitN-GFP-R | ATTTGCGGCCGCATTCAAGGGCGGATTTGAC | | (*EcoR* I and *Not* I sites are underlined) |
| DUF-GFP-F | GGGGTACCCAATACTCCAAGAAATGGC | 882 | Expression of DUF-GFP in S2 cells |
| DUF-GFP-R | ATTTGCGGCCGCAGGTGTAAGGGTAAGATC | | (*Kpn* I and *Not* I sites are underlined) |
| vWD-GFP-F | CGGAATTCTGCGTTGCTGACAAGATGC | 609 | Expression of vWD-GFP in S2 cells |
| vWD-GFP-R | ATTTGCGGCCGCAACAGTTTTGAGGGGCGGT | | (*EcoR* I and *Not* I sites are underlined) |
| TYLCV CP-GST-F | CGGGATCCATGTCGAAGCGACCAGGCGA | 793 | Expression of TYLCV CP in *E. coli* |
| TYLCV CP-GST-R | CGGAATTCTTAATTTGATATTGAATCAT | | (*BamH* I and *EcoR* I sites are underlined) |

*T7, 5´-TAATACGACTCACTATAGG- 3´
